# Characterisation of Posterior Predominant Amyloid PET Binding Across Multiple Cohorts

**DOI:** 10.64898/2026.06.17.733031

**Authors:** Joseph Giorgio, Ganna Blazhenets, Susan M Landau, Stefania Pezzoli, Jennifer S. Yokoyama, David N. Soleimani-Meigooni, Maria C Carrillo, Lea T Grinberg, William W Seeley, Salvatore Spina, Kelly N Nudelman, Liana G Apostolova, Bradford C Dickerson, William J Jagust, Gil D Rabinovici, Renaud La Joie, The LEADS Consortium, Alzheimer’s Disease Neuroimaging Initiative

**Author notes:** Correspondence: Joseph Giorgio, Department of Biostatistics, University of California Berkeley, Berkeley, California, USA. Data used in preparation of this article were obtained from the Alzheimer’s Disease Neuroimaging Initiative (ADNI) database (adni.loni.usc.edu). As such, the investigators within the ADNI contributed to the design and implementation of ADNI and/or provided data but did not participate in analysis or writing of this report. A complete listing of ADNI investigators can be found at: http://adni.loni.usc.edu/wpcontent/uploads/how_to_apply/ADNI_Acknowledgement_List.pdf.

## Abstract

The standard approach to quantify amyloid (Aβ) PET averages uptake within a single cortical mask that assumes no clinically relevant spatial heterogeneity in uptake patterns. Here, in a sample of 12,379 clinically impaired participants taken from four phenotypically diverse cohorts we use data-driven approaches to discover heterogeneous patterns of Aβ-PET binding, uncovering a reproducible and clinically relevant pattern of posterior predominant Aβ-PET binding. In particular, Aβ-PET positive participants who have posterior predominant binding are less likely to be *APOE*-ε4 carriers, more severely impaired, have thinner cortex in posterior regions, and greater posterior tau PET burden. Furthermore, in a subsample of participants with neuropathological assessment, participants with posterior predominant Aβ binding have a higher likelihood of having cerebral amyloid angiopathy at autopsy. These findings suggest that Aβ-PET accumulates along two orthogonal axes with biological and clinical relevance, indicating that the standard approach to assess Aβ-PET is insufficient to capture meaningful signal from Aβ-PET imaging. This work has implications for the diagnosis and treatment of Alzheimer’s disease and extends our understanding of the mechanisms governing variable Aβ-PET distribution and its downstream effects.

## Introduction

The deposition of cerebral amyloid-beta (Aβ) assessed via Aβ-PET imaging is the gold standard for in vivo diagnosis of Alzheimer’s Disease (AD). To quantitatively assess Aβ-PET load, the standard approach derives a single scalar value by averaging binding across regions in the frontal, temporal, and parietal lobes (i.e. centiloid CL) ^1^. Similarly, the FDA-approved guidelines for visually appraising Aβ-PET positivity in clinical settings rely on the detection of tracer binding in a single cortical area, without interpreting regional patterns^2^. Whether a single global scalar adequately captures the pathologically relevant information in Aβ-PET, or if there is clinically relevant signal in the distribution of Aβ-PET signal remains to be determined.

The Thal neuropathological staging scheme describes a sequential progression of Aβ plaques across the brain, starting with neocortical deposition in the association cortex, without specifying regional involvement (phase 1), followed by involvement of medial temporal regions (phase 2), striatum (phase 3), midbrain (phase 4), and finally involvement of the cerebellum (phase 5)^3^. Consistent with this scheme, the progressive involvement of neocortical followed by striatal Aβ-PET signal has been shown to track with disease severity^4^. When investigating the trajectory of neocortical Aβ-PET accumulation, staging studies have put forward a single trajectory, with midline anterior and posterior regions showing abnormality first and occipital and somatosensory regions showing abnormality in later stages ^5–9^. It is unclear if this neocortical trajectory fully captures the heterogeneity in Aβ-PET binding and accounts for all relationships between Aβ-PET with clinical variability and other AD pathologies.

Prior work has associated elevated occipital Aβ-PET binding to cerebral amyloid angiopathy (CAA) ^10–17^, a neuropathologically diagnosed condition where Aβ deposits in the cerebrovascular wall that is frequently co-morbid with AD ^18^. In addition, work in small samples has reported subtle increases in posterior Aβ-PET binding in AD patients with Posterior Cortical Atrophy (PCA) compared to other AD phenotypes ^19,20^, although this finding has been inconsistent ^21^. Whether data driven approaches to differentiate global and regional Aβ-PET effects support Aβ-PET imaging phenotypes for these conditions is not clear and requires substantially larger sample sizes than previous analyses.

Here, we apply data-driven approaches to voxel-wise Aβ-PET imaging to explore whether distinct, reproducible patterns of Aβ-PET binding exist in clinically impaired, Aβ-PET positive participants. Using a large real-world clinic-based sample from the Imaging Dementia-Evidence for Amyloid Scanning (IDEAS) study (n=10,361) we built a model to characterise spatial heterogeneity in Aβ-PET binding. We then applied data from three independent and clinically diverse cohorts (n=2,018) to this model to determine the replicability of any identified patterns and their associations with neuropathology, clinical, neuroimaging, and genetic markers for AD. By building a model on highly heterogenous clinical scans and applying it to clinically diverse external data we extend prior subtyping work ^22,23^ to provide an in-depth and robust investigation of data driven Aβ-PET phenotypes and their clinical and biological relevance.

## Results

### Participants

We aggregated data from four cohorts of clinically impaired participants with a clinical diagnosis of mild cognitive impairment (MCI) or mild dementia resulting in a total sample of 12,379 patients. These participants had varying degrees of multimodal data available with all having at least one Aβ-PET session (**Table 1**). We used the IDEAS (n=10,361) study to develop our imaging phenotype discovery model and perform preliminary group wise contrasts. IDEAS Aβ-PET was acquired using either [^18^F]-Florbetaben (n=3033), [^18^F]-Florbetapir (n=6699) or [^18^F]-Flutemetamol (n=629) and these Aβ-PET images were the only input into the discovery model. We applied Aβ-PET from three out-of-sample cohorts to the model fit on IDEAS data. Out-of-sample datasets of participants with MCI or dementia included the [^18^F]-Florbetaben Aβ-PET from the Longitudinal Early-Onset Alzheimer’s Disease Study (LEADS n=509) a multicentre study of sporadic early onset AD, the [^18^F]-Florbetaben (n=177) or [^18^F]-Florbetapir Aβ-PET (n=816) from the Alzheimer’s Disease Neuroimaging Initiative (ADNI n= 993) a multicentre study of late onset amnestic AD, and the [^11^C]-Pittsburgh compound B Aβ-PET from the University of California, San Francisco Alzheimer’s Disease Research Center (UCSF n=516) a clinical cohort including patients with clinical syndromes associated with AD or frontotemporal lobar degeneration (FTLD). These deeply phenotyped datasets were used as out-of-sample cohorts for associating different Aβ-PET phenotypes with multiple AD biomarkers and neuropathological assessment.

**Table 1.** Sample characteristics. MMSE: Mini-Mental State Examination; FBB: Florbetaben; FBP: Florbetapir; FLUTE: Flutemetamol; PiB: Pittsburgh Compound B; FTLD: frontotemporal lobar degeneration

|  | Primary cohort<br>– model<br>development | Secondary cohorts – external validation |  |  |
| --- | --- | --- | --- | --- |
| Cohort | IDEAS | LEADS | ADNI | UCSF |
| **General description** | multicentre real-world study of Aβ-PET in patients > 65yo | multicentre study of sporadic early onset AD | multicentre study of late onset amnesic AD | single site study of patients with FTLN- and AD-associated syndromes |
| **Sample size** | 10,361 | 509 | 993 | 516 |
| **Age mean ± std** | 75.7±6.3 | 58.9±4.6 | 73.9±8.1 | 64.4±9.4 |
| **Female Sex (%F)** | 5245 (51%) | 256 (51%) | 434(44%) | 246(48%) |
| **MCI/Dementia (%Dem)** | 6500/3861 (37%) | 173/332 (65%) | 736/256 (26%) | 317/199 (38%) |
| **MMSE mean ± std** | 24.4±5 | 22.4±5.4 | 26.6±3.2 | 23.3±4.2 |
| **Centiloids mean ± std** | 45±49.1 | 79±49.1 | 49.3±50.3 | 50±51.4 |
| **Aβ Tracer (FBB/FBP/FLUTE/PiB)** | 3033/6699/629/0 | 509/0/0/0 | 177/816/0/0 | 0/0/0/516 |
| **APOE-ε4 (0/1/2) alleles (%ε4 carriers)** | 568/471/114 (51%) | 244/188/63 (51%) | 454/346/104 (50%) | 291/150/36 (39%) |
| **Flortaucipir-PET (n)** | n.a | 482 | 431 | 301 |
| **Autopsy (n)** | n.a | n.a | 49 | 166 |

### A**β**-PET phenotypes

Within the IDEAS sample we used spatial independent components analysis (ICA) to extract components (n dimensions=40) of grey matter Aβ-PET uptake. Through visual inspection of the 40 components, we retained 11 ICA components that showed clear grey matter binding excluding remaining components capturing non-relevant signal (e.g. white matter, ventricles, skull, etc.) (**Supplementary Figure 1**). Next, we used k-means clustering on each participant’s grey matter scores (i.e. participant level scalar representing Aβ-PET uptake in each component). We found the first partition of the sample (i.e. k=2) separated participants based on Aβ-PET positivity, the second partition (i.e. k=3) split the Aβ positive group into two groups. When partitioning the sample again (i.e. k=4) one of the Aβ positive groups was almost entirely preserved (**Supplementary Figure 2**). Based on the aim of interrogating differences in Aβ positive groups, and the consistent assignment of participants to the same cluster at lower and higher dimensionality (i.e. k=3 and k=4) we chose 2 partitions (i.e. k=3) to cluster participants, resulting in two Aβ positive clusters and one Aβ negative cluster.

Visualising the centroid locations for each of the three clusters we observed low binding across the cortex in the Aβ negative cluster (IDEAS: n=4729, mean CL=2), and within the Aβ positive clusters: the Aβ+_(typical)_ (IDEAS: n=3148, mean CL=86) had loading in components capturing the canonical regions used for Aβ-PET quantification (i.e. CL like) whereas the Aβ+_(posterior)_ (IDEAS: n=2484, mean CL=76) cluster had binding across the cortex with predominant binding in occipital components (**Supplementary Figure 3)**. We ran post-hoc evaluations on these data to further understand the different binding patterns between the Aβ+_(posterior)_ and Aβ+_(typical)_ groups. First, we projected each participant’s Aβ-PET scan into the two leading principal components of grey matter binding, then we projected the k-means centroid locations onto this basis set. Visualising the data in these two dimensions we observed two primary axes of variation: an Aβ positivity axis explaining 36% of the variance that separates the Aβ positive clusters from the Aβ negative cluster and a second axis explaining 20% of the variance in grey matter binding that separates the Aβ+_(posterior)_ and Aβ+_(typical)_ clusters (**Figure 1a)**. To better understand the variable binding patterns related to these clusters we calculated the average Aβ-PET image for participants assigned to these clusters (**Figure 1b**) and subtracted these from each other, revealing a posterior-anterior gradient in Aβ-PET binding (**Figure 1c**.). This gradient shows the Aβ+_(posterior)_ population is not more burdened globally than the Aβ+_(typical)_ group, but rather the Aβ+_(posterior)_ group has markedly less Aβ-PET uptake in the frontal lobes. Calculating these mean difference maps for each tracer independently produced the same gradient, indicating this was not a feature of Aβ-PET tracer (**Supplementary Figure 4**).

**Figure 1.**
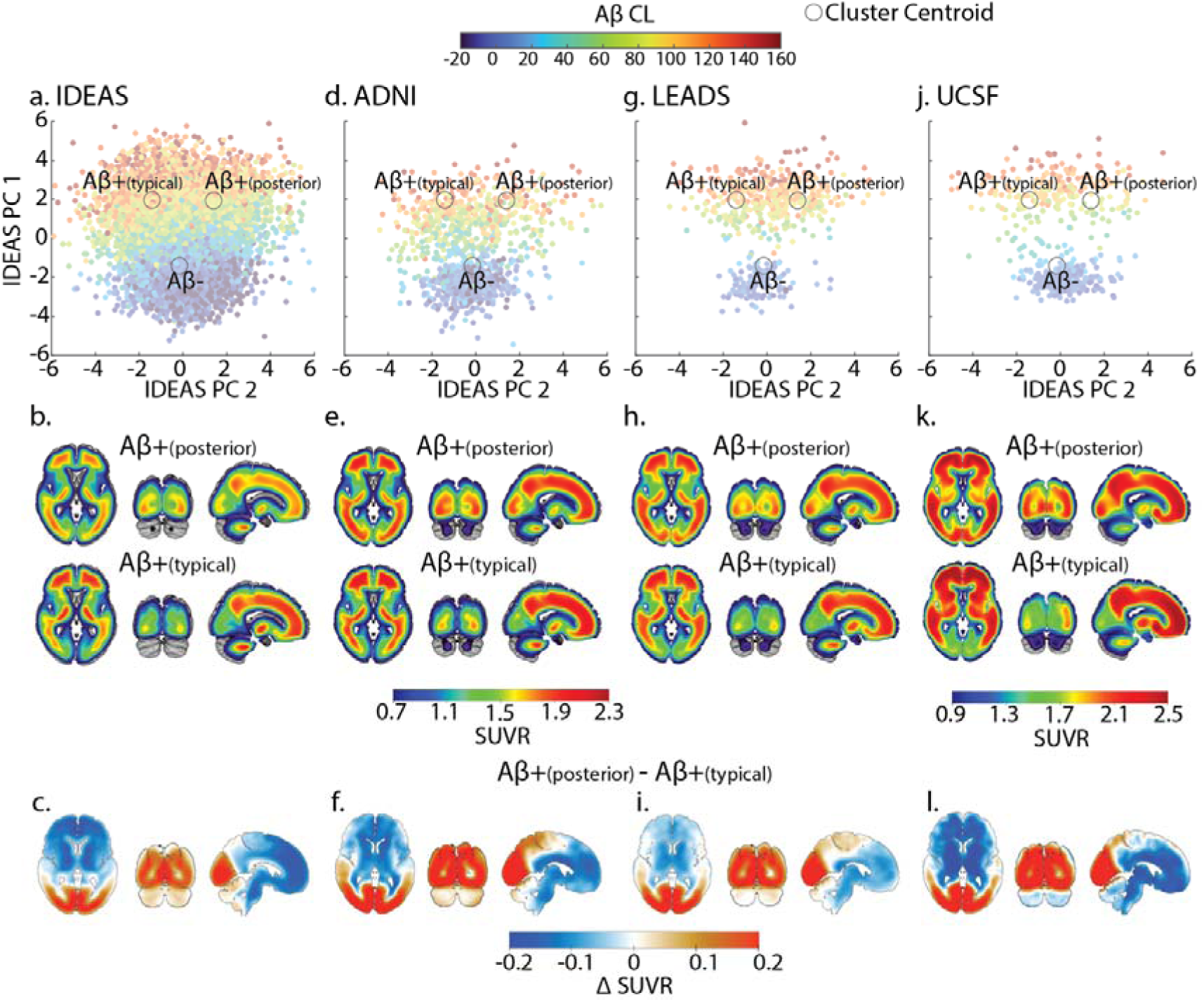
model derived posterior. **A**β**-PET phenotype.** Scatter plots represent the Aβ-PET scans for each participant in a. IDEAS, d. ADNI, g. LEADS, j. UCSF projected onto the first two leading dimensions of grey matter binding as defined using the IDEAS sample only. The colour of each point represents the centiloid (CL) value for that scan. Black circles are the centroid locations defined using k-means clustering on the IDEAS sample projected onto the first two dimensions of grey matter binding. Average whole brain standardised uptake value ratio (SUVR) from the Aβ-PET scans from the patients classified as Aβ+_(posterior)_ or Aβ+_(typical)_ b. IDEAS, e. ADNI, h. LEADS, k. UCSF are projected on the MNI brain. UCSF Aβ-PET scans were collected using [^11^C]-PiB and average uptake images (k) are shown on a different scale to the cohorts that utilise [^18^F] tracers. Difference between average Aβ-PET SUVR for Aβ+_(posterior)_ vs. Aβ+_(typical)_ participants from c. IDEAS, f. ADNI, i. LEADS, l. UCSF independent cohorts.

Applying out-of-sample data to the model fit on the IDEAS data replicated the posterior-anterior gradient in the other three datasets (**Figure 1)**. Furthermore, across the four independent samples variance along the Aβ+_(typical)_ to Aβ+_(posterior)_ axis increased with higher values on the Aβ positivity axis. This suggests that moving along the Aβ+_(typical)_ to Aβ+_(posterior)_ gradient is not directly coupled with greater levels of Aβ-PET, rather it indicates that the model has identified two additive processes, one related to Aβ positivity and another relating to variance in the posterior-anterior topography of uptake in Aβ-PET positive participants.

In general, there was a higher proportion of Aβ+ participants assigned to the Aβ+_(typical)_ cluster compared to the Aβ+_(posterior)_ cluster, except for the LEADS cohort where proportions were similar (% Aβ+_(posterior)_ vs. Aβ+_(typical)_: IDEAS 24% vs. 30%; ADNI 20% vs. 28%; UCSF 20% vs. 25%; LEADS 38% vs. 38%) (**Supplementary Figures 5a**). Contrasting CL values for each Aβ positive cluster showed no consistent differences in overall Aβ load as defined within standard regions for quantification (**Supplementary Figure 5b**).

We next used longitudinal Aβ-PET in the ADNI (n participants=497; n scans=1316; inter-scan interval= mean 2.25±std 0.85 years) and LEADS (n participants =295; n scans=788; inter-scan interval= mean 1.33±std 0.6 years) samples to assess how within-subject assignment to the Aβ-PET clusters transitioned over time. So as not to bias the analysis by number of scanning sessions per participant we calculated the transitions between all adjacent scans, pooling data within and across participants in each cohort to determine transition probabilities (e.g. a participant with four scans will have three transitions included in the calculation of transition probability). We observed that cluster assignment over time was consistent (**Supplementary Figure 6**), with both cohorts showing similar within cluster stability ranging from 85-95%. For the few between cluster transitions, 10.5% (ADNI) and 13.5% (LEADS) of early observations of Aβ+_(typical)_ transitioned to Aβ+_(posterior)_ scans at a later timepoint. Whereas 4.5% (ADNI) and 1.3% (LEADS) of early observations of Aβ+_(posterior)_ transitioned to Aβ+_(typical)_ scans at a later timepoint. Finally, we observed in participants assigned to the Aβ negative cluster at one observation there are similar proportions moving to either the Aβ+_(posterior)_ (4% ADNI; 6.1% LEADS) or the Aβ+_(typical)_ (3.5% ADNI; 9.1% LEADS) cluster indicating divergent pathways from Aβ negative to either Aβ positive group (**Supplementary Figure 6**). Although participants are more likely to transition from Aβ+_(typical)_ to Aβ+_(posterior)_ than vice versa, the divergent pathways from Aβ negative to either Aβ+_(posterior)_ or Aβ+_(typical),_ suggests that there is not a single canonical pathway from Aβ negative to Aβ+_(typical)_ to Aβ+_(posterior)_.

### Demographic and clinical features of A**β**-PET phenotypes

We next compared sex, age, and MMSE for each Aβ cluster (**Supplementary Figure 7**). As our main goal was to characterise the demographic and clinical correlates of Aβ-PET patterns rather than differences between Aβ negative and Aβ positive individuals, which has been reported on extensively^24–27^, we omitted the Aβ negative cluster from further analyses. We observed no consistent age differences between the Aβ+_(posterior)_ vs. Aβ+_(typical)_ groups (**Supplementary Figure 7b)**. However, there were reproducible sex differences between the two Aβ+ groups, with the Aβ+_(posterior)_ cluster having a significantly (p<0.05) higher frequency of females in three out of four cohorts (%female Aβ+_(posterior) vs._ Aβ+_(typical),_ IDEAS: 55% vs. 50%; LEADS: 59% vs. 48%; ADNI: 41% vs. 43% (not significant); UCSF: 62% vs. 47%) (**Supplementary Figure 7a)**. We consistently observed that the Aβ+_(posterior)_ group included significantly fewer *APOE*-ε4 carriers than the Aβ+_(typical)_ group (no *APOE*-ε4 alleles Aβ+_(posterior)_ vs. Aβ+_(typical),_ IDEAS: 38% vs. 28%; LEADS: 53% vs. 36%; ADNI: 34% vs. 23%; UCSF: 55% vs. 38) (**Figure 2**).

**Figure 2.**
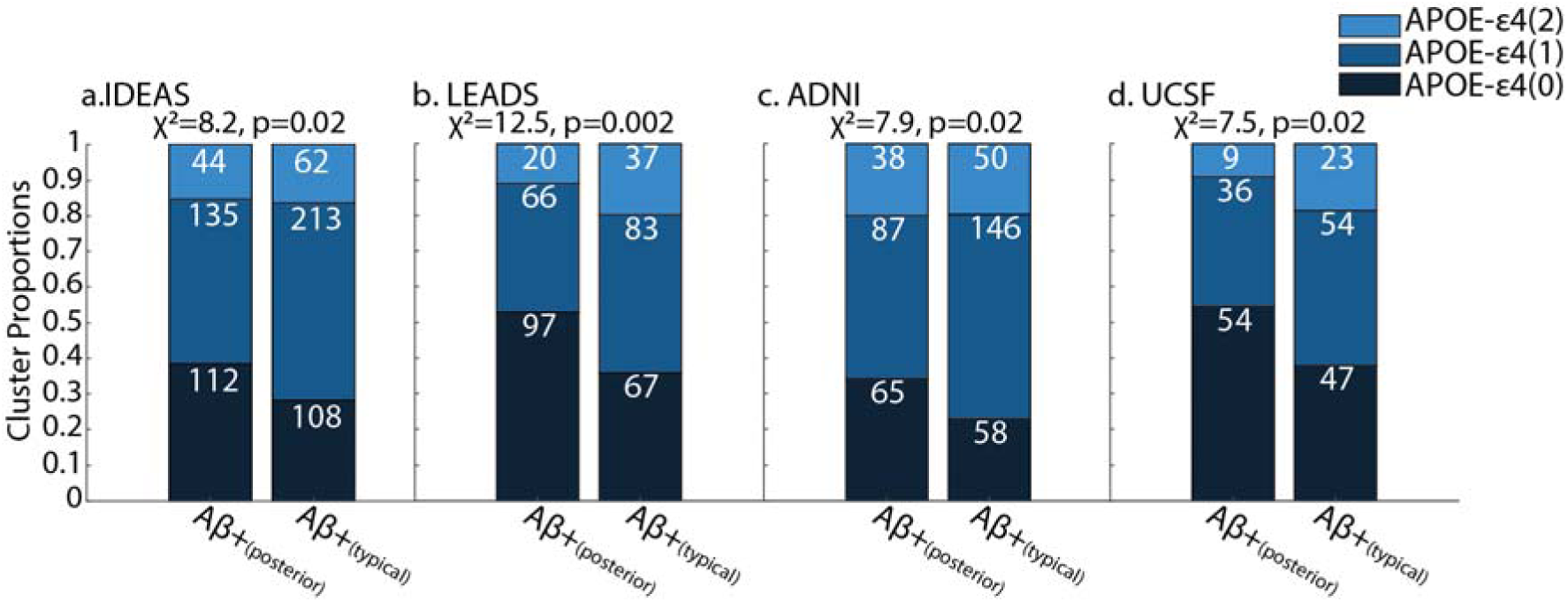
**differences in number of *APOE*-**ε**4 alleles between A**β**+_(posterior)_ and A**β**+_(typical)_ groups.** Stacked barcharts show the proportion of groups with 0, 1, or 2 *APOE*-ε4 alleles in Aβ+_(posterior)_ and Aβ+_(typical)_ groups for each cohort. Values in white are the counts for each grouping based on number of *APOE*-ε4 alleles. Chi square statistics are shown above each barchart indicating differences in the proportions of *APOE*-ε4 dosage between the Aβ+_(posterior)_ and Aβ+_(typical)_ groups in each cohort.

There were marginal and weak group differences in MMSE, with the Aβ+_(posterior)_ cluster having significantly lower MMSE score in all cohorts except the UCSF sample (mean MMSE Aβ+_(posterior)_ vs. Aβ+_(typical)_: IDEAS 23.3 vs. 23.6, Cohen’s d =-0.069; LEADS 20.2 vs. 22.4, Cohen’s d =-0.404; ADNI 24.8 vs. 25.8 Cohen’s d =-0.31; UCSF 21.0 vs. 21.5, Cohen’s d =-0.068 (not significant)) (**Supplementary Figure 7c)**. When comparing clinical diagnosis of MCI or dementia however, participants assigned to the Aβ+_(posterior)_ cluster were more clinically impaired in all cohorts (%Dementia Aβ+_(posterior)_ vs. Aβ+_(typical),_ IDEAS: 47% vs. 41%; LEADS: 82% vs. 64%; ADNI: 48% vs. 35%; UCSF: 54% vs. 39%) (**Figure 3**).

**Figure 3.**
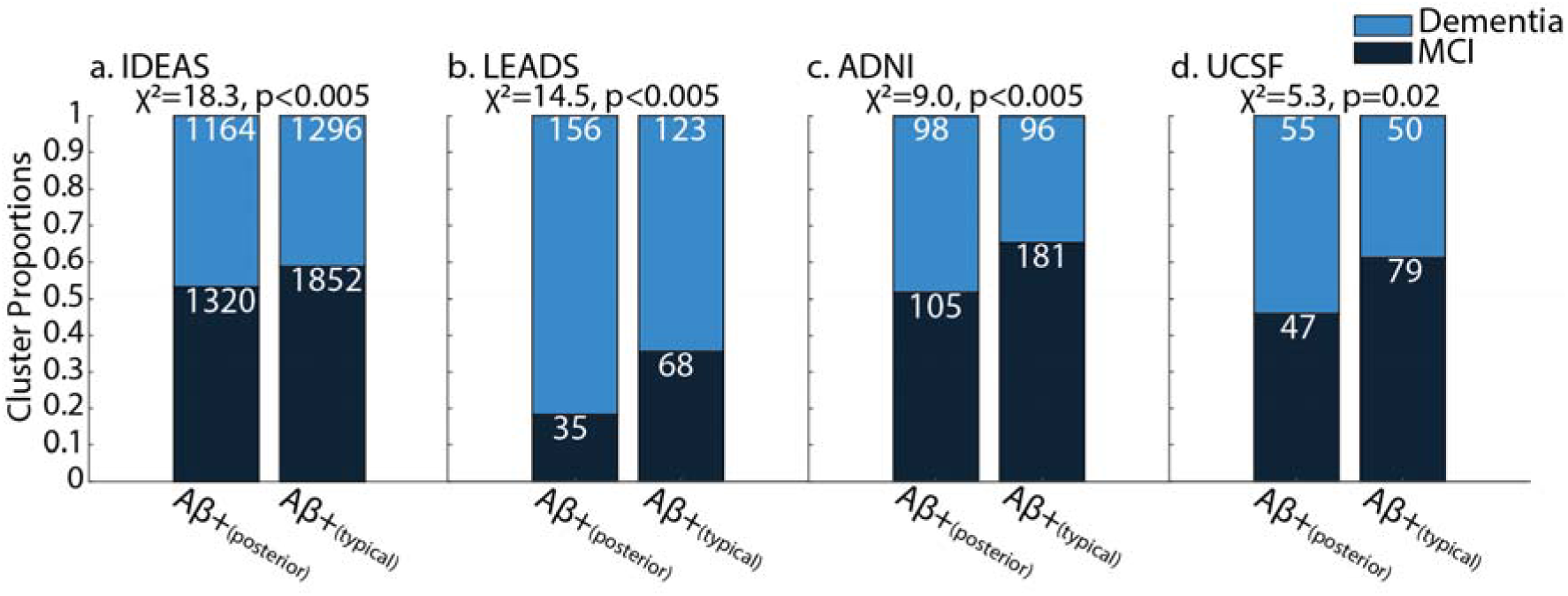
differences in cognition between. **A**β**+_(posterior)_ and A**β**+_(typical)_ groups.** a. differences in clinical impairment between Aβ+_(posterior)_ and Aβ+_(typical)_ groups in each cohort. Chi square statistics are shown above each barchart indicating significant differences in the proportions of clinical severity between the Aβ+_(posterior)_ and Aβ+_(typical)_ groups in each cohort.

Contrasting different cognitive domain scores in the ADNI and LEADS samples revealed a consistent trend showing relatively larger deficits in executive function domains in participants assigned to Aβ+_(posterior)_ compared to Aβ+_(typical)_. In particular the Aβ+_(posterior)_ group reliably showed greater deficits in executive and processing speed functions than the Aβ+_(typical)_ cluster (ADNI Executive Function t(439)=-4.55, p<0.001, Cohen’s d =-0.437; LEADS Working Memory t(294)=-5.36, p<0.001, Cohen’s d =-0.623 / Speed and Attention: t(294)=-4.34, p<0.001, Cohen’s d =-0.516) (**Figure 4**). Within the phenotypically heterogeneous LEADS and UCSF samples we contrasted the proportions of participants with Aβ+_(posterior)_ vs. Aβ+_(typical)_ within a given clinical phenotype. We observed amnestic participants were less likely to have an Aβ+_(posterior)_ binding (amnestic participants LEADS: Aβ+_(posterior)_ 45.6% χ^2^=11.3, p<0.001; UCSF: Aβ+_(posterior)_ 38.5%, χ^2^=3.9, p=0.048) whereas participants with a clinical diagnosis of PCA were more likely to have an Aβ+_(posterior)_ binding (PCA participants LEADS: Aβ+_(posterior)_ 77% χ^2^=8.2, p=0.004; UCSF: Aβ+_(posterior)_ 68%, χ^2^=6.5, p=0.011) (**Figure 4**).

**Figure 4.**
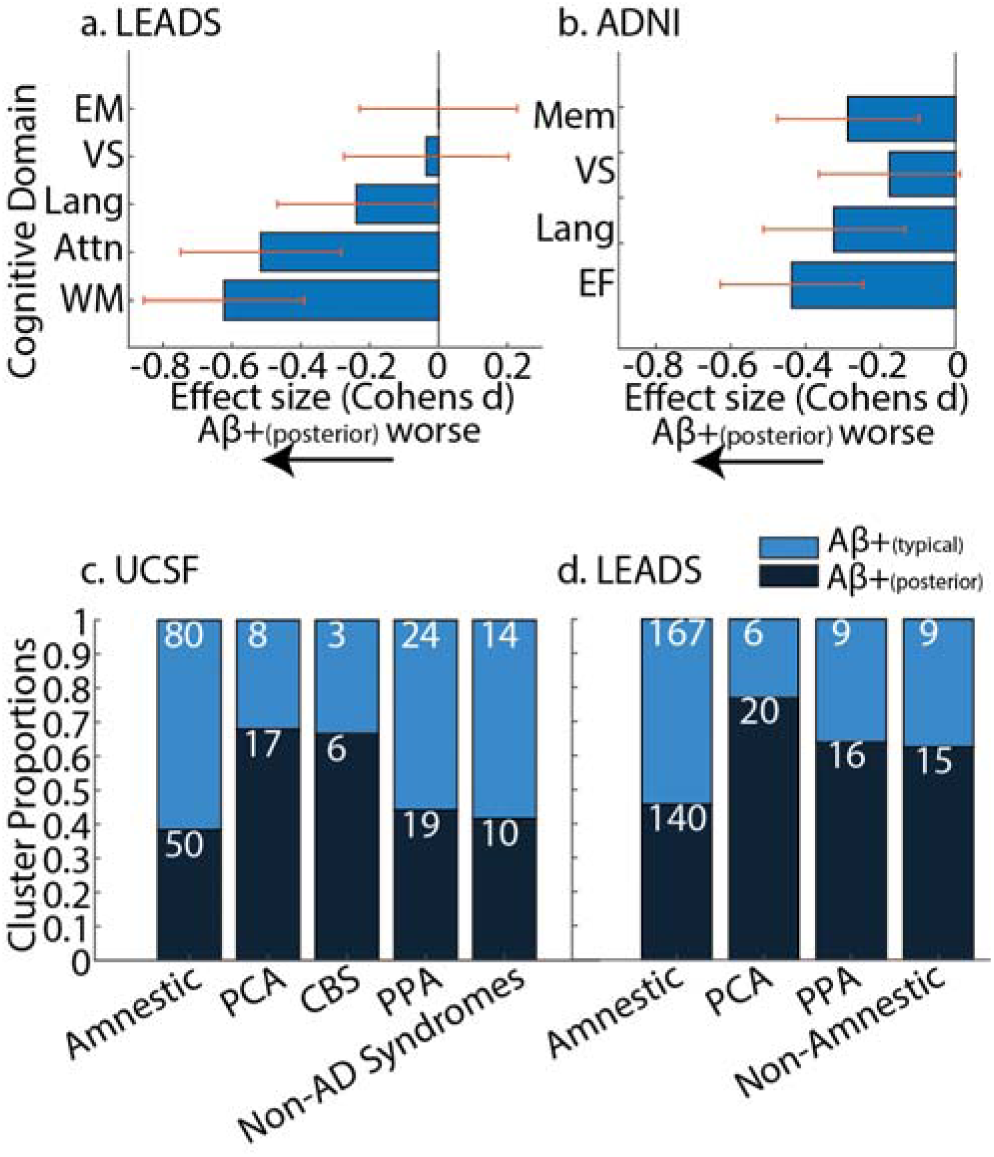
differences in cognition between. **A**β**+_(posterior)_ and A**β**+_(typical)_ groups.** Differences in cognitive domains between Aβ+_(posterior)_ and Aβ+_(typical)_ groups in the a. LEADS and b.

ADNI cohort. Barcharts show the effect size of the difference (two sample t-test) between neuropsychological composites, red whiskers show the 95% confidence interval of the effect size estimate. A negative value represents a lower (worse) score in that cognitive domain for the Aβ+_(posterior)_ group. Differences in the proportion of participants with Aβ+_(posterior)_ and Aβ+_(typical)_ PET binding in different clinical phenotypes in the c. UCSF and d. LEADS cohorts. Dementia syndromes are amnestic, Posterior Cortical Atrophy (PCA), Corticobasal syndrome (CBS), Primary Progressive Aphasia (PPA), other non-AD, or non-amnestic AD syndromes. Stacked barcharts represent the proportion of each group, where the white number shows the number of patients represented in each segment.

### Associations of A**β**-PET phenotypes with other imaging modalities

Next, we contrasted other in vivo imaging markers between the two Aβ positive clusters within the three secondary cohorts; these analyses could not be run in IDEAS as the study did not include imaging data beyond Aβ-PET. White matter hyperintensity volumes were available for both ADNI and LEADS cohorts, showing higher lesion volume in the Aβ+_(posterior)_ vs Aβ+_(typical)_ groups (ADNI t(433)=2.232, p=0.03, Cohen’s d=0.216; LEADS t(359)=3.392, p<0.005, Cohen’s d=0.361) (**Figure 5a,b**). Contrasting regional cortical thickness revealed consistent differences, with the Aβ+_(posterior)_ group having lower cortical thickness in occipital, parietal, and posterior temporal lobes (**Figure 5c**). To exclude the possibility that Aβ positive PET differences were an artifact due to head size we compared total intracranial volume observing no differences (**Supplementary Figure 8a.**). Finally, we assessed if there were differences in bilateral hippocampal volume between the two Aβ positive clusters with only LEADS showing significant differences indicating the Aβ+_(posterior)_cluster has greater bilateral hippocampal volume (Aβ+_(posterior)_ vs. Aβ+_(typical)_ LEADS t(368)=2.664, p=0.008) (**Supplementary Figure 8b.**). Finally, tau-PET was available in the three secondary cohorts and consistently revealed greater posterior tau-PET uptake in the Aβ+_(posterior)_ group compared to the Aβ+_(typical)_ group (**Figure 5d**.).

**Figure 5.**
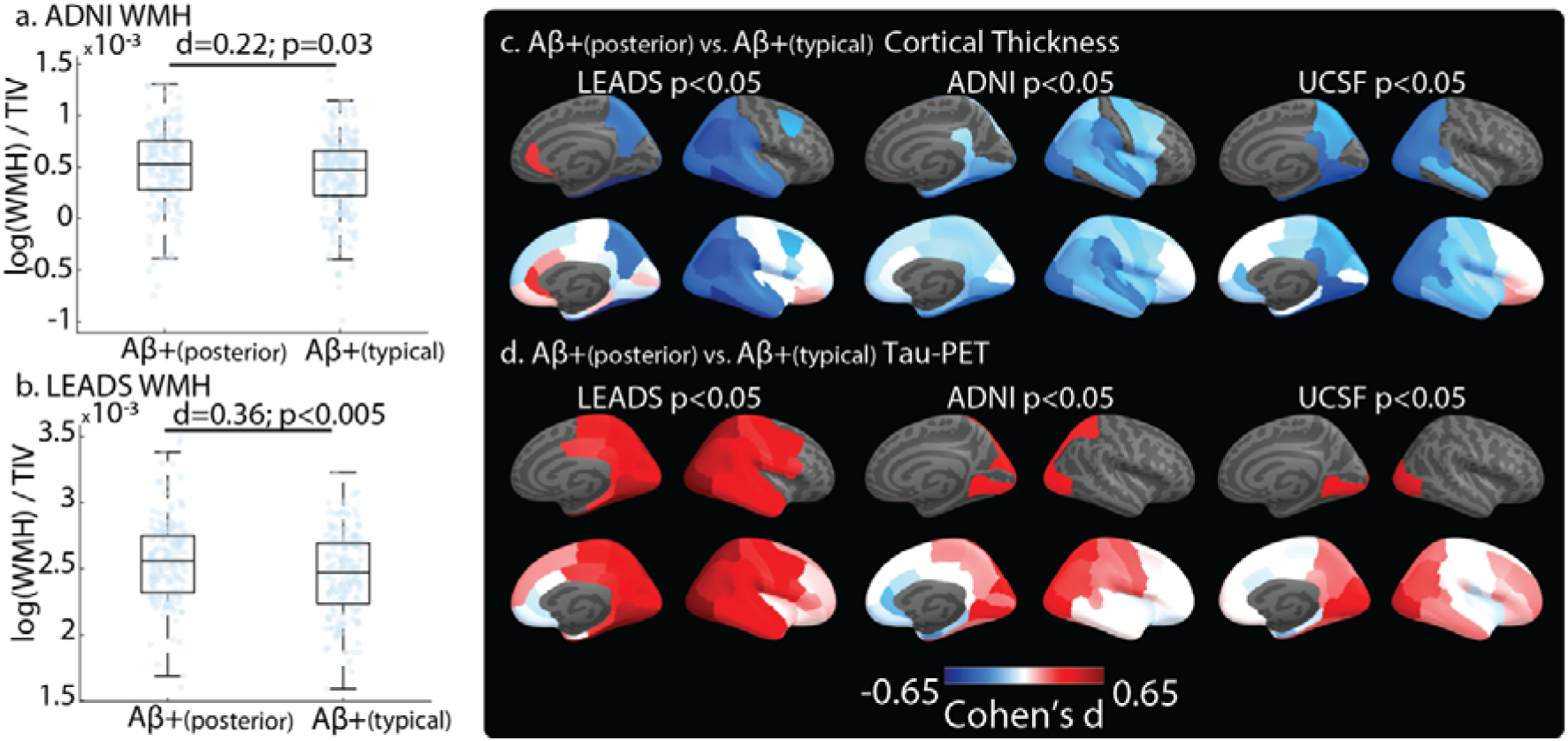
differences in tau-PET, MRI thickness and white matter hyperintensities between. **A**β**+_(posterior)_ and A**β**+_(typical)_ groups.** Differences in normalised volumes of white matter hyperintensities (WMH) between Aβ+_(posterior)_ and Aβ+_(typical)_ groups in a. ADNI and b. LEADS. Effect size (Cohen’s d) and significance value for the two-sample t-test comparing WMH intensities is shown above the boxplots. Differences in averages of left and right hemisphere c. cortical thickness and d. tau PET uptake measured in 34 ROIs from the Desikan-Killiany atlas between Aβ+_(posterior)_ and Aβ+_(typical)_ groups. ROIs shown in blue indicate a lower a. cortical thickness (more pathological) and b. lower tau PET uptake (less pathological) in the Aβ+_(posterior)_ population, whereas areas in red indicate higher a. cortical thickness (less pathological) and b. higher tau-PET (more pathological) uptake. Top panels show regions with significant differences (p<0.05 two sample t-test), bottom panel shows the effect sizes (Cohen’s d) for the differences between Aβ+_(posterior)_ and Aβ+_(typical)_ groups across the cortex.

### Associations of A**β**-PET phenotypes with neuropathology

Finally, postmortem assessment was available in a subset of the UCSF and ADNI participants (n=215 total; **Supplementary Table 1**). Due to limited samples in each cohort, particularly after excluding the 100 participants assigned to the Aβ negative cluster, we collapsed UCSF and ADNI datasets into a single sample resulting in 116 participants who were either Aβ+_(posterior)_ or Aβ+_(typical)_ ante-mortem. We regressed cluster membership (Aβ+_(posterior)_ or Aβ+_(typical)_) on the degree of CAA pathology (none, mild, moderate, severe) at autopsy to determine if the topography of Aβ-PET predicted more severe CAA. We observed participants with an Aβ+_(posterior)_ pattern of Aβ-PET binding presented with more severe CAA pathology at autopsy (t(114)=2.45 β=0.14, p=0.015) (**Figure 6**), with consistent relationships within each cohort (**Supplementary Figure 9**). Comparing the proportion of participants with any CAA in the combined UCSF and ADNI sample, we observed that 93% of Aβ+_(posterior)_ have evidence of any CAA compared to 79% of individuals with Aβ+_(typical)_ Aβ-PET binding (χ^2^=4.56, p=0.033). In general, Aβ+_(typical)_ patients tended to be at a more advanced Braak stage than Aβ+_(posterior)_ patients (t(114)=1.93; β=0.08, p=0.056) (**Figure 6**), with consistent relationships within each cohort (**Supplementary Figure 9 a,b.**). Variability in the stage of tau pathology is limited in this sample with most participants showing advanced tau pathology (i.e. Braak 5/6) reducing statistical power. Finally, we did not observe any clear differences in Aβ histopathological stage, with most participants having Thal stage 4/5 pathology (**Supplementary Figure 10 a,b.**). Similarly, there did not appear to be clear differences in the presence or distribution of Lewy body pathology (**Supplementary Figure 10 e,f.**).

**Figure 6.**
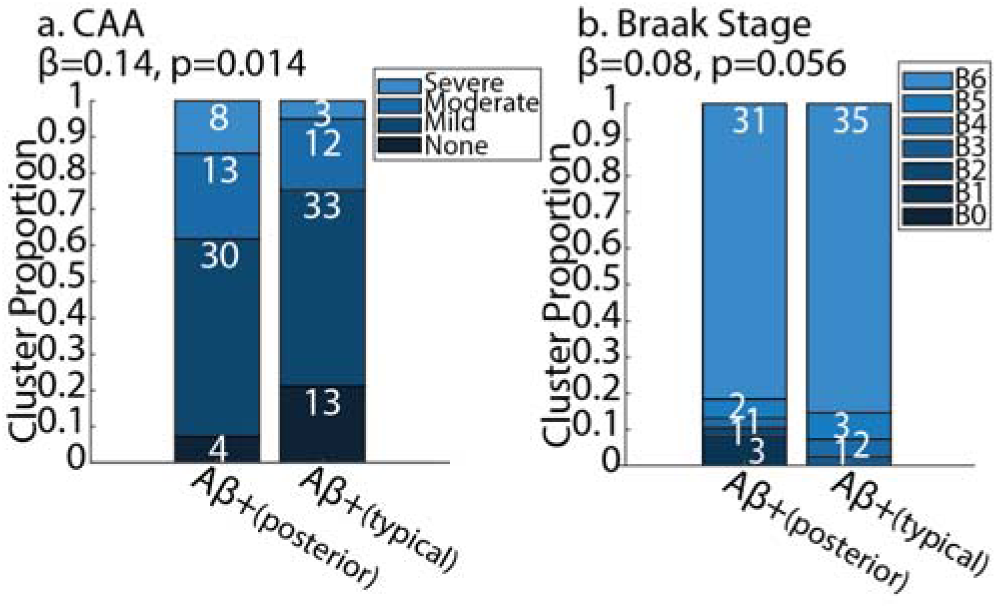
Comparison of histologically assessed cerebral amyloid angiopathy and tau severity between Aβ+_(posterior)_ and Aβ+_(typical)_ groups in ADNI and UCSF. a. severity of cerebral amyloid angiopathy (CAA). b. severity of tau pathology assessed using Braak staging. Stacked barcharts represent the proportion of each group, where the white number shows the number of participants represented in each segment. Sample is comprised of UCSF and ADNI participants who went to autopsy with antemortem Aβ positive PET. The values above each barchart show the parameter estimate (β) and significance (p value) when regressing each participant’s ante-mortem Aβ-PET cluster on the severity of a. CAA (t(114)=2.45 β=0.14, p=0.015), or b. Braak Stage (t(114)=1.9; β=0.08, p=0.056).

## Discussion

Here, we provide a detailed exposition of posterior-predominant Aβ-PET binding. Using a purely data-driven approach we find a reproducible posterior to anterior gradient of Aβ-PET binding in clinically impaired participants who are Aβ-PET positive. We observe robust associations between other imaging modalities in three heterogeneous out-of-sample cohorts, highlighting those participants with a posterior predominance of Aβ-PET harbour more tau and thinner cortex in posterior cortical regions. These participants are more clinically impaired, more likely to present with PCA, and present with greater cognitive impairment in executive functions, while also being less likely to carry the *APOE*-ε4 allele. This work represents the first in-depth exploration of posterior Aβ-PET binding and has implications for understanding regional susceptibility to tau, atrophy, and how this may lead to specific cognitive deficits and clinical phenotypes.

Using latent modelling and clustering of whole brain Aβ-PET we uncovered three reliable clusters of participants, with two divergent patterns in Aβ positive participants. These Aβ positive participants had either an Aβ+_(typical)_ pattern capturing regions used in CL quantification and a more Aβ+_(posterior)_ group that has decreased frontal but -at a similar magnitude- increased occipital binding, a region omitted from CL quantification. We observed in participants from LEADS and ADNI who converted from Aβ negative to Aβ positive at a later scanning session, there was a similar likelihood of moving to either Aβ+_(posterior)_ or Aβ+_(typical)_ suggesting divergent pathways towards Aβ-PET positivity. Most previous in vivo Aβ-PET studies have investigated a single trajectory of Aβ-PET abnormality; with midline anterior and posterior regions showing abnormality first and occipital and somatosensory regions showing abnormality in later stages, with these later stages associated with greater current and incipient clinical impairment ^5,6,8,9,22,28–31^. Our findings suggest these single-trajectory models may not be adequate to capture heterogeneity in Aβ-PET binding. Rather, we propose there are two additive processes involved in the topography of Aβ-PET binding, one related to Aβ positivity and another relating to the posterior-anterior topography of Aβ-PET uptake. This finding is supported in recent clustering and subtyping analyses showing the existence of an occipital predominant Aβ-PET binding pattern ^22,23^.

We show that participants with posterior predominant Aβ-PET binding are more likely to have evidence of CAA at autopsy, with just 7% showing no histological evidence of CAA (compared to 21% of Aβ+_(typical)_ participants). Furthermore, we observed that participants classified as Aβ+_(posterior)_ have a higher load of white matter hyperintensities, a correlate of CAA ^32^. Differential binding patterns of Aβ-PET have been reported in CAA with patients showing an increased relative binding in occipital regions ^10–17^. These previous studies investigated small sample sizes, however when meta-analysed, an occipital: global ratio did separate CAA compared to AD ^33^, a measure that was subsequently validated ^34^. These previous studies however were limited in that they did not generally assess neuropathologically confirmed CAA as an outcome, rather they investigated radiographic assessment of CAA using MRI. A study investigating the relationship between regional PiB PET with CAA at autopsy showed no differences in occipital Aβ-PET binding relative to CAA burden^35^. Here, we have shown that in AD participants (i.e. Aβ-PET positive) that neuropathologically confirmed CAA is associated with differences in the topography of Aβ-PET, with Aβ+_(posterior)_ participants highly likely to present with CAA and with greater severity of CAA. Having an Aβ+_(posterior)_ binding pattern is a sensitive marker for CAA, however it is not a specific marker of CAA; as 79% of patients with Aβ+_(typical)_ Aβ-PET binding still had some level of CAA at autopsy. As such, it appears an Aβ+_(posterior)_ pattern may be useful to rule in CAA, but an Aβ+_(typical)_ pattern cannot rule out CAA.

We observe a clinical, neuropsychological, and neuroimaging phenotype associated with posterior predominant Aβ-PET binding. Aβ+_(posterior)_ participants presented with thinner posterior cortices, greater posterior tau-PET burden, and greater clinical impairment showing more pronounced deficits in functions supported by neocortical regions (e.g. executive function). However, these Aβ+_(posterior)_ patients have similar, or in the case of LEADS, greater hippocampal volume and less frontal Aβ-PET burden. These dissociable findings suggest that presenting with Aβ+_(posterior)_ binding is not just an advanced stage of sporadic AD but rather presents a more focal posterior phenotype of AD in contrast to a more typical amnestic AD phenotype (i.e. Aβ+_(typical)_).

Greater thinning in posterior regions of the cortex has been shown in patients with posterior predominant Aβ-PET binding^36^, along with more severe cognitive impairment ^22,23,30,36,37^.

However, the spatial concordance between posterior predominant Aβ-PET binding and tau-PET binding shown here has not been widely reported. These findings were reproduced in three independent cohorts with effect sizes in a moderate range and are consistent with our phenotypic results showing that 77% of LEADS and 68% of UCSF PCA patients have a posterior predominant Aβ-PET binding pattern. The link between PCA and neuroimaging findings in occipito-parietal and occipito-temporal cortex is well reported on, showing increases in tau^38^ and hypometabolism^39^, and decreases in synaptic density^40^ and MRI grey matter thickness^41^. Here, in two relatively large independent samples of PCA patients who were Aβ-PET positive, we observe an association between posterior predominant Aβ-PET and PCA, supporting previous findings in considerably smaller samples ^19,20^. However, even in non PCA syndromes, patients classified as Aβ+_(posterior)_ still present with the tau-PET and MRI hallmarks of PCA. Together, this suggests the posterior predominant imaging phenotype may not be confined to PCA as a discrete syndrome but rather may reflect a broader spectrum of posterior cortical change potentially initiated by greater posterior Aβ burden.

Within the two Aβ-PET positive subgroups we observed reproducible differences in *APOE*-ε4 carriage rate with an association between the presence of the *APOE*-ε4 allele and Aβ+_(typical)_ PET binding. Alternative data driven approaches to separate Aβ-PET imaging similarly described a subgroup with more posterior binding pattern ^22,23^, with both studies observing an association with more anterior Aβ-PET binding and carriage of the *APOE*-ε4 allele. Furthermore, using univariate associations a relative reduction in occipital Aβ-PET binding (i.e. more anterior binding) was associated with a higher proportion of *APOE*-ε4 carriers^36^. This robust association between more anterior Aβ positive PET binding and a higher proportion of *APOE*-ε4 carriers occurred across four independent samples presented here, as well as one from South Korea^36^, the Mayo Clinic ^22^, and a merged sample of six cohorts^23^. *APOE*-ε4 has been routinely linked to a reduced homeostatic clearing of Aβ ^42–44^. Furthermore, *APOE* is not uniformly expressed across the cortex, with gene expression showing an anterior to posterior gradient (**Supplementary Figure 11**). Considering this, it is plausible that the increased prevalence of *APOE*-ε4 carriership in the Aβ+_(typical)_ group may be indicative of a selective vulnerability of the frontal lobes to Aβ through impaired Aβ clearance due to the maladaptive APOE4 isoform.

Finally, in three of the four cohorts, women were more likely to present with Aβ+_(posterior)_ binding than men. Prior work in the IDEAS dataset has shown women are more likely to be read as visually Aβ-PET positive and have on average higher CL levels than men^45^.

However, it appears women are also more likely to have a posterior predominant Aβ-PET binding, a pattern that is associated with greater clinical severity. When considered alongside evidence that women have high medial temporal tau burden for a given level of Aβ-PET than men^46^, this represents an additional layer of pathophysiological burden on women that may further explain the higher prevalence of AD dementia in women^47^. This novel finding warrants further exploration to further understand the complex relationships between AD pathologies, sex, and other biological processes.

There are several limitations to this work. First, despite the Aβ-PET imaging sample being very large the subsequent sub analyses involved substantially lower numbers. In particular, the Aβ-PET positive antemortem sample consisted of only 116 individuals who were at advanced stages of AD limiting the range of Thal and Braak staging. Caution is also required when directly comparing Aβ-PET imaging to Thal and Braak stages, as these neuropathological schemes provide a qualitative rather than quantitative assessment of regional Aβ and tau pathology. Apparent discordance between Aβ-PET topography and neuropathological staging may also reflect sensitivity limitations of Aβ-PET, which requires a threshold of Aβ plaque density before binding becomes detectable. Additional work incorporating quantitative neuropathology is needed to more firmly ground the Aβ-PET imaging phenotypes to the underlying distribution and level of Aβ plaques. Furthermore, when investigating the association between Aβ-PET phenotypes and CAA neuropathology CAA was modelled as none, mild, moderate, severe following NACC criteria. This approach does not capture the staging of CAA with respect to the anatomical compartment of deposition, whereby leptomeningeal involvement precedes parenchymal involvement, and leptomeningeal CAA is known to show a posterior predominance^48^. A more fine-grained appraisal of regional CAA with quantitative measurements is required to fully characterise its relationship to posterior Aβ-PET binding. Second, the longitudinal assessment of Aβ-PET cluster over time was limited and only assessed in clinically impaired individuals. Further work covering a longer evolution of Aβ-PET from lower to higher levels is required to better understand how different Aβ-PET phenotypes develop over time. Third, the interrogation of cognitive domains was limited to only the ADNI and LEADS samples each with limitations on the neuropsychological composites available. The ADNI memory composite does not separate out episodic and working memory, with the latter more implicated in executive functioning rather than mnemonic function predominantly served by medial temporal lobes. Furthermore, there are limited visuospatial tests administered in ADNI and therefore this composite may not be sensitive to visuospatial deficits. With regards to the visuospatial domain, it is likely that in the LEADS sample there are censoring and floor effects on these tests that limit its sensitivity to detect visuospatial deficits in impaired participants.

The findings presented here have several potential implications for the treatment of AD as well as expanding our understanding of clinical variability in AD and its association to the topography of Aβ-PET. First, we observed that the presentation of an Aβ+_(posterior)_ topography is related to a high likelihood of having neuropathologically confirmed CAA. CAA is a major risk factor for Amyloid Related Imaging Abnormalities (ARIA), a common side effect of approved monoclonal antibodies for the removal of Aβ plaques ^49,50^. As such, the topography of Aβ-PET in Aβ positive patients may be relevant when treating and managing patients on anti-amyloid therapies. Second, the omission of the occipital lobes in the standard centiloid quantification or the visual reading of Aβ-PET scan regardless of tracer underestimates the true Aβ-PET burden in a scan. This exclusion of the occipital lobe also represents a bias that underestimates the explainable heterogeneity in clinical severity based on the topography of Aβ-PET, with a relative increase in posterior binding compared to frontal binding reliably explaining variance in clinical severity. Third, it has been thought that there is limited information in the spatial distribution of Aβ-PET with little to no association with clinical or other imaging variables. However, we see that there are reproducible relationships between the topography of Aβ-PET, cortical thinning, and tau-PET uptake, and that this posterior predominance in pathology is associated with deficits in cognitive functioning supported by more neocortical regions (i.e. non-amnestic). Fourth, we observed that there is a reliable effect of *APOE*-ε4 carriage on the distribution of Aβ-PET providing a possible link between the anterior to posterior gradient of *APOE* gene expression and the distribution of Aβ-PET. However, *APOE* is not the only gene expressed in an anterior to posterior gradient and it is feasible that there are polygenic effects of genes expressed along this gradient that may govern the distribution of cortical Aβ. This provides a new avenue to understand the underlying drivers of the topography of pathologies along the AD cascade that warrants further investigation.

## Conclusions

Aβ-PET is the gold standard for in vivo quantification of cortical Aβ and diagnosis of AD. Despite the proliferation of this tool in research and clinical contexts the meaning of spatial variability in Aβ-PET binding has been incompletely characterised. Here we provide evidence in clinically impaired, Aβ-PET positive participants, for a robust association between posterior predominant Aβ-PET binding and more severe clinical impairment, posterior atrophy, and posterior tau burden. Together, this suggests the current convention of aggregating Aβ-PET positive, clinically impaired individuals without regard for Aβ-PET topography warrants reconsideration.

## Methods

### Participants

We sampled participants with MCI or dementia from four independent cohorts (n= 12,379) who underwent Aβ-PET imaging. Data from the IDEAS study (n=10,361, mean age±std= 75.7±6.3, MCI/Dementia =6500/3861) was used for model formulation. IDEAS was a multicentre real-world study of the clinical utility of Aβ-PET imaging within a clinical setting^51^. IDEAS participants were over 65 years old and were diagnosed with either MCI or dementia prior to Aβ-PET imaging. Three external datasets with extensive multimodal data available were then applied to the model fit on IDEAS data. The LEADS study (n=509, mean age±std= 58.9±4.6, MCI/Dementia =173/332) is a multicentre study of sporadic early onset AD, participants presented with varied phenotypes including amnestic, non-amnestic, posterior cortical atrophy and primary progressive aphasia^52^. Participants had either an MCI or dementia diagnosis and were less than 65 years old. The UCSF (n=516, mean age±std= 64.4±9.4, MCI/Dementia =317/199) dataset includes participants diagnosed with clinical syndromes associated with either frontotemporal lobar degeneration or AD, including AD dementia, corticobasal syndrome, posterior cortical atrophy, primary progressive aphasia, progressive supranuclear palsy, behavioural variant frontotemporal dementia, as well as patients with mixed pathologies or Parkinson’s disease dementia. ADNI (n=993, mean age±std= 73.9±8.1, MCI/Dementia =736/256) is a multicentre study of late onset amnestic AD and includes participants with MCI or dementia who present with predominantly amnestic deficits. The ADNI was launched in 2003 as a public-private partnership, led by Principal Investigator Michael W. Weiner, MD. The primary goal of ADNI has been to test whether serial magnetic resonance imaging (MRI), positron emission tomography (PET), other biological markers, and clinical and neuropsychological assessment can be combined to measure the progression of mild cognitive impairment (MCI) and early Alzheimer’s disease (AD).

### A**β**-PET imaging

*IDEAS:* IDEAS participants were scanned using either [^18^F]-Florbetaben (n=3033), [^18^F]-Florbetapir (n=6699) or [^18^F]-Flutemetamol (n=629). Aβ-PET images were processed using rPOP^53^ (https://github.com/leoiacca/rPOP), an MRI free processing pipeline resulting in an image in MNI template space smoothed to a standard resolution of 10mm^3^ and resliced to 2mm^3^ ([101,116,96]) voxel dimensions. Voxel values were normalised to SUVR using the whole cerebellum as a reference region. SUVR images then underwent a visual quality control process.

*LEADS:* LEADS participants had Aβ PET scans acquired using [^18^F]-Florbetaben. Tracer was administered via intravenous injection (∼8 mCi), followed by acquisition of 4 × 5 minute PET frames at 90-110 minutes post-injection.

*ADNI:* ADNI participants were scanned using either [^18^F]-Florbetaben (n=177) or [^18^F]-Florbetapir (n=816). The [^18^F]-Florbetapir Aβ-PET protocol involved injection of 10 mCi of [^18^F]-Florbetapir followed by the acquisition of 20 min of emission data at 50-70 min post injection. The [^18^F]-Florbetaben protocol involved injection of 8.1 mCi of [^18^F]-Florbetaben followed by 20min of emission data at 90-110min post injection.

*UCSF:* UCSF participants were scanned using [^11^C]-Pittsburgh compound B. PET data were acquired from 0-90 min (35 frames) or 50-70 min (4 x 5 min frames) after the injection of ∼15 mCi of [^11^C]-Pittsburgh Compound-B. Data from a 50-70 min post-injection acquisition window was selected to calculate SUVR images.

For ADNI, LEADS, and UCSF Aβ-PET frames were realigned, averaged, and co-registered onto their corresponding MRI. SUVR images were created in native space using an MRI-defined whole cerebellar reference region. These SUVR images were then warped to MNI space and smoothed to a resolution of 10mm^3^ and resliced to 2mm^3^ ([101,116,96]) voxel dimensions. Centiloid values were derived from these values following cohort specific pipelines and transformation equations that have been described previously (IDEAS ^54^; LEADS ^52^; ADNI ^55^; UCSF^56^)

### MRI

Structural MRIs for UCSF participants were acquired at UCSF, on a 3T Siemens Tim Trio or a 3T Siemens Prisma Fit scanner. Both scanners had similar acquisition parameters (sagittal slice orientation; slice thickness = 1 mm; slices per slab = 160; in-plane resolution = 1 x 1 mm; matrix = 240x256; repetition time = 2.3 ms; inversion time = 900 ms; flip angle = 9°; echo time: Trio: 2.98 ms; Prisma: 2.9 ms). LEADS and ADNI structural T1 weighted MRIs are acquired using similar sequences based on the ADNI protocol. Images were a sagittal 3D accelerated MPRAGE/IRSPGR T1-weighted sequence: TR/TE/TI = 2300/3/900 ms, flip angle 9°, sagittal orientation, FOV = 256 × 240 mm with 208 slices, 1 × 1 × 1 mm resolution, and 2x acceleration. All Structural T1-weighted images were analysed using similar FreeSurfer based processing pipelines. Images were first segmented into grey matter, white matter, and CSF components in native space using Statistical Parametric Mapping (SPM12). Native space T1 images were additionally parcellated with FreeSurfer v.7 using the Desikan-Killiany atlas. Cortical thickness in these regions of interest (ROIs) was extracted as well as volume of the hippocampus and total intracranial volume. In addition to the T1 weighted MRI, fluid-attenuated inversion recovery (FLAIR) sequences were acquired and segmented to measure the volume of white matter hyperintensities (WMH) following previously published methods in LEADS^57^ and ADNI^58^ .

### Tau-PET imaging

A subset of ADNI (n=431), LEADS(n=482) and UCSF (n=301) participants were scanned with [^18^F]-Flortaucipir tau PET using a similar protocol. The [^18^F]-Flortaucipir-PET protocol entailed the injection of 10 mCi of FTP followed by acquisition of 30 min of emission data from 75-105 min post injection. [^18^F]-Flortaucipir-PET frames were realigned, averaged, and co-registered onto their corresponding MRI. SUVR images were created in native space using an MRI-defined inferior cerebellar grey matter reference region. Using the FreeSurfer ROIs from the corresponding T1 MRI the average [^18^F]-Flortaucipir-PET uptake in bilateral ROIs was extracted.

### Post-mortem neuropathological assessment

Participants in ADNI (n=49) and UCSF (n=166) with ante-mortem Aβ-PET underwent histological assessment of the cortex for Aβ plaques, tau tangles, neuritic plaques, Lewy bodies, and cerebral amyloid angiopathy^59,60^. The severity of AD neuropathology was assessed using Braak staging for tau, Thal phases for Aβ deposition, and CERAD scoring for neuritic plaque density (none, sparse, moderate, frequent). The severity of cerebral amyloid angiopathy was quantified as none, mild, moderate, and severe. Lewy bodies were assessed based on their presence in the brain stem, limbic regions, neocortical regions, olfactory bulb, or amygdala.

### Cognitive assessment

All participants underwent neuropsychological assessment. In the IDEAS study a diagnosis of MCI or dementia was made by a dementia specialist within the past 24 months. In addition, most participants underwent mental status testing with the Mini-Mental State Examination (MMSE). In ADNI, clinical diagnosis of MCI or dementia was performed based on the Petersen criteria^61^. Participants underwent extensive neuropsychological assessment including the MMSE; neuropsychological measures were used to derive composites that covered the memory ^62^, executive functioning^63^, language, and visuospatial domains^64^.

LEADS diagnosis was performed through multidisciplinary consensus conference at each clinical site following the NIA-AA diagnostic criteria for dementia and MCI; in addition a diagnosis of logopenic aphasia and posterior cortical atrophy were made using published criteria^65,66^. Extensive neuropsychological assessment was performed in LEADS including the NACC Uniform Data Set cognitive battery, the NACC Frontotemporal Lobar Degeneration module, the Alzheimer’s Disease Cooperative Studies - cognitive behaviour subscale (ADAS Cog), and several additional cognitive tests tapping into cognitive functions that are commonly impaired in rare AD variants ^52^. This neuropsychological data was then used to derive cognitive composites covering episodic memory, working memory, speed and attention, language, and visuospatial domains. The UCSF diagnosis was performed through multidisciplinary clinical consensus conferences involving neurologists and neuropsychologists, integrating neurological examination, informant interview, and comprehensive cognitive testing. Participants underwent an extensive neuropsychological assessment that included standardized batteries (including NACC Uniform Data Set measures and additional domain-specific tests) assessing memory, executive function, language, and visuospatial abilities, alongside functional and behavioural evaluations. Diagnoses of MCI and dementia were assigned based on established clinical criteria following review of all available clinical and cognitive data.

### *APOE*-ε4 genotyping

The number of *APOE*-ε4 alleles was assessed in each cohort with varying levels of participant coverage (IDEAS n= 1153; LEADS n= 495; ADNI n=904; UCSF n= 477). In the UCSF sample, *APOE*-ε4 genotype was assigned from the rs429358 and rs7412 SNPs defining the epsilon locus using a TaqMan assay^67^. For ADNI *APOE*-ε4 genotype was downloaded from LONI. In LEADS, following DNA extraction at National Centralized Repository for Alzheimer’s Disease and Related Dementias (NCRAD), genotyping was performed using a custom 96-SNP fingerprint panel that includes assays for rs429358 and rs7412 ^52^. In IDEAS, *APOE*-ε4 genotype was obtained through the Amyloid Neuroimaging and Genetics Initiative add-on study, with DNA derived from saliva samples banked at NCRAD.

### A**β**-PET Phenotype discovery

*Spatial ICA*: To reduce the high dimensionality and treat for sources of noise in the Aβ-PET images we apply group spatial ICA (i.e. source based morphometry) to the IDEAS SUVR images in MNI space. We used the sbm module in the GIFT toolbox (https://trendscenter.org/software/gift/) to perform the ICA. Initial pre-processing involved masking the Aβ-PET scan with a binary mask encompassing CSF, grey matter and white matter in the MNI152 template brain, removing the mean voxel value for each Aβ-PET scan within the masked region, and principal component analysis (40 PCs). ICA was then run (40 ICs) using the Infomax algorithm with spatial maps generated using back reconstruction. The result of this analysis is a set of brain maps of voxel loadings for each component, ICA_Loadings_ 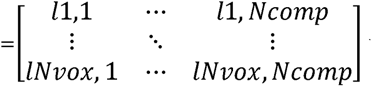 where *l* is the loading on a voxel *Nvox* is the number of voxels in the Aβ-PET scans and *Ncomp* is the number of independent components. From this matrix and an Aβ-PET scan a score for each independent component is derived for IDEAS participant representing the multivariate weighted sum of the Aβ-PET load within each component 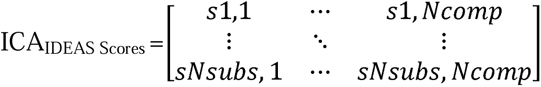 where *s* is the score for a component and *Nsubs* is the number of IDEAS subjects. From the 40 estimated components we performed visual inspection of the brain loading maps to determine which components captured grey matter signal and which were noise. We selected 40 components to provide sufficient dimensionality to separate structured grey matter signal from noise. Although minimum description length criterion estimated a two-dimensional solution reflecting the dominant positive and negative binding patterns, this was insufficient to resolve the range of noise present in PET data. A 20-component solution produced highly similar downstream cluster assignments, supporting the robustness of the clustering to ICA dimensionality and the suitability of this approach for characterising the dominant axis of variation in grey matter Aβ-PET binding.

*k-means clustering*: Next we determined if there were groupings of participants based on the spatial topography of their Aβ-PET binding. To do this we performed k-means clustering on each participant’s scores (ICA_IDEAS Scores_) for components showing grey matter binding. Prior to running the clustering we z-scored the component scores using the mean (µ_ICA IDEAS Scores_) and std (σ _ICA IDEAS Scores_) of the IDEAS sample: Norm ICA_IDEAS Scores =_ (ICA_IDEAS Scores–_ µ_ICA IDEAS Scores_)/ σ _ICA IDEAS Scores_, we retained the scaling vectors µ_ICA IDEAS Scores_ and σ _ICA IDEAS Scores_ for rescaling external datasets. On the Norm ICA_IDEAS Scores_ matrix we ran k-means using a squared Euclidean distance metric, running the clustering for k= 2,3, or 4 clusters. To select the number of clusters (k) of participants we generated an alluvial diagram tracking the flow of participants membership as the dimensionality increased and qualitatively assessing if different partitions of the sample were a parent to higher dimensional partitions (i.e. if multiple high dimensional clusters were nested in the same lower dimensional cluster). We favoured this to standard approaches for assessing the number of clusters (i.e. silhouette, gap or elbow approaches) as these all converge on a bimodal solution (k=2) separating Aβ positive and Aβ negative participants.

The result of this analysis is the assignment of each IDEAS patient to a cluster based on the topography of their Aβ-PET binding and the co-ordinates of the centroids in the normalised ICA scores space, 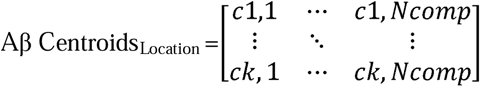 where *c* is the co-ordinate of the centroid for each ICA grey matter component, indicating the relative feature importance for each cluster. We perform post-hoc investigation of the clustering by running a principal component analysis on the grey matter scores derived from the ICA. Using the PCA coefficients we then project the cluster centroids and each IDEAS participants grey matter scores into the two leading PCs. The result of this analysis is a coefficient matrix, PCA_Loadings_ 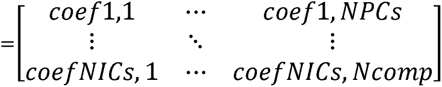 that we retain to project new data into the IDEAS principal component subspace.

*Applying new data to IDEAS model:* Using the model parameters learnt on the IDEAS data we apply several simple linear operations to assign new patients to a cluster of Aβ-PET binding. With the exception of the UCSF sample we use IDEAS scaling parameters to deterministically assign each participant to a given cluster (i.e. independent of out-of-sample cohort distributions of Aβ-PET pathology). As the UCSF sample uses [^11^C]-Pittsburgh Compound-B which has a different dynamic range than the IDEAS [^18^F] tracers we re-calculate the component means and standard deviations in the UCSF sample when normalising subject scores as a z-scaled matrix of data for cluster assignment.

To assign new data to the model built on IDEAS data, we project a given participants (*Sub*) Aβ-PET scan onto the ICA grey matter scores basis set.

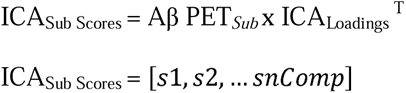

We next normalise the participants scores using the scaling parameters defined in the IDEAS sample.

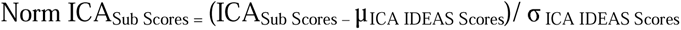

For UCSF participants who were scanned using [^11^C]-Pittsburgh Compound-B we perform a different within sample scaling.

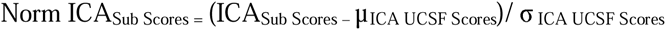

Next, we determine the minimum squared Euclidean distance from the subjects Norm ICA_Sub Scores_ to the cluster centroids learnt in the IDEAS sample and assign subject (*Sub*) the closest cluster.

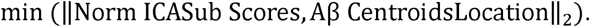

To perform diagnostics on how well the new data fits into the subspace learnt on IDEAS data we project the subjects Norm ICA_Sub Scores_ into the PCA basis set learnt on IDEAS data

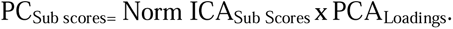

### Software and data availability

ICA, clustering and statistical analysis was run in MATLAB 2023b using the statistics and machine learning toolbox, GIFT and SPM12. All code and learnt matrices to project new Aβ-PET data into the ICA and PCA basis set, as well as perform cluster assignment is provided in the supplementary information of this publication. Software to perform pre-processing of the Aβ-PET data is available at (https://github.com/leoiacca/rPOP). Data access can be requested through each independent research study (UCSF: https://memory.ucsf.edu/research-trials/professional/open-science; LEADS: https://leads-study.medicine.iu.edu/researchers/leads-data-request-application/; IDEAS: https://www.ideas-study.org/Original-Study/Data-Request; ADNI: https://ida.loni.usc.edu/collaboration/access/appLicense.jsp).

## Acknowledgements

J.G. is supported by the Alzheimer’s Association (23AARF-1026883). Data collection and sharing for this project was funded by the Alzheimer’s Disease Neuroimaging Initiative (ADNI) (National Institutes of Health Grant U01 AG024904) and DOD ADNI (Department of Defense award number W81XWH-12-2-0012). ADNI is funded by the National Institute on Aging, the National Institute of Biomedical Imaging and Bioengineering, and through generous contributions from the following: AbbVie, Alzheimer’s Association; Alzheimer’s Drug Discovery Foundation; Araclon Biotech; BioClinica, Inc.; Biogen; Bristol-Myers Squibb Company; CereSpir, Inc.; Cogstate; Eisai Inc.; Elan Pharmaceuticals, Inc.; Eli Lilly and Company; EuroImmun; F. Hoffmann-La Roche Ltd and its affiliated company Genentech, Inc.; Fujirebio; GE Healthcare; IXICO Ltd.; Janssen Alzheimer Immunotherapy Research & Development, LLC.; Johnson & Johnson Pharmaceutical Research & Development LLC.; Lumosity; Lundbeck; Merck & Co., Inc.; Meso Scale Diagnostics, LLC.; NeuroRx Research; Neurotrack Technologies; Novartis Pharmaceuticals Corporation; Pfizer Inc.; Piramal Imaging; Servier; Takeda Pharmaceutical Company; and Transition Therapeutics. The Canadian Institutes of Health Research is providing funds to support ADNI clinical sites in Canada. Private sector contributions are facilitated by the Foundation for the National Institutes of Health (www.fnih.org). The grantee organization is the Northern California Institute for Research and Education, and the study is coordinated by the Alzheimer’s Therapeutic Research Institute at the University of Southern California. ADNI data are disseminated by the Laboratory for Neuro Imaging at the University of Southern California. The IDEAS study was funded by the Alzheimer’s Association, the American College of Radiology, Avid Radiopharmaceuticals Inc. (a wholly owned subsidiary of Eli Lilly and Company), General Electric Healthcare, and Lantheus.

LEADS is funded by the National Institute on Aging (NIA) U01-AG057195 and NIA R56-AG057195. Work performed in the UCSF ADRC and affiliated studies was supported by NIH/NIA P30-AG062422, R35-AG072362, R01-NS050915, P01-AG019724 and the Rainwater Charitable Foundation. J.S.Y. received funding from NIH-NIA R01AG062588, R01AG057234, and U19AG079774; NIH-NINDS U54NS123985; the Rainwater Charitable Foundation; the Alzheimer’s Association; the Global Brain Health Institute; and the Mary Oakley Foundation. The content of this publication is solely the responsibility of the authors and does not necessarily represent the official views of the NIH. Lantheus and Avid Radiopharmaceuticals enabled the use of florbetaben and flortaucipir but did not provide direct funding and were not involved in data analysis or interpretation.

