## Supplementary for "Characterisation of Posterior Predominant Amyloid PET Binding Across Multiple Cohorts"

**Supplementary Figures**

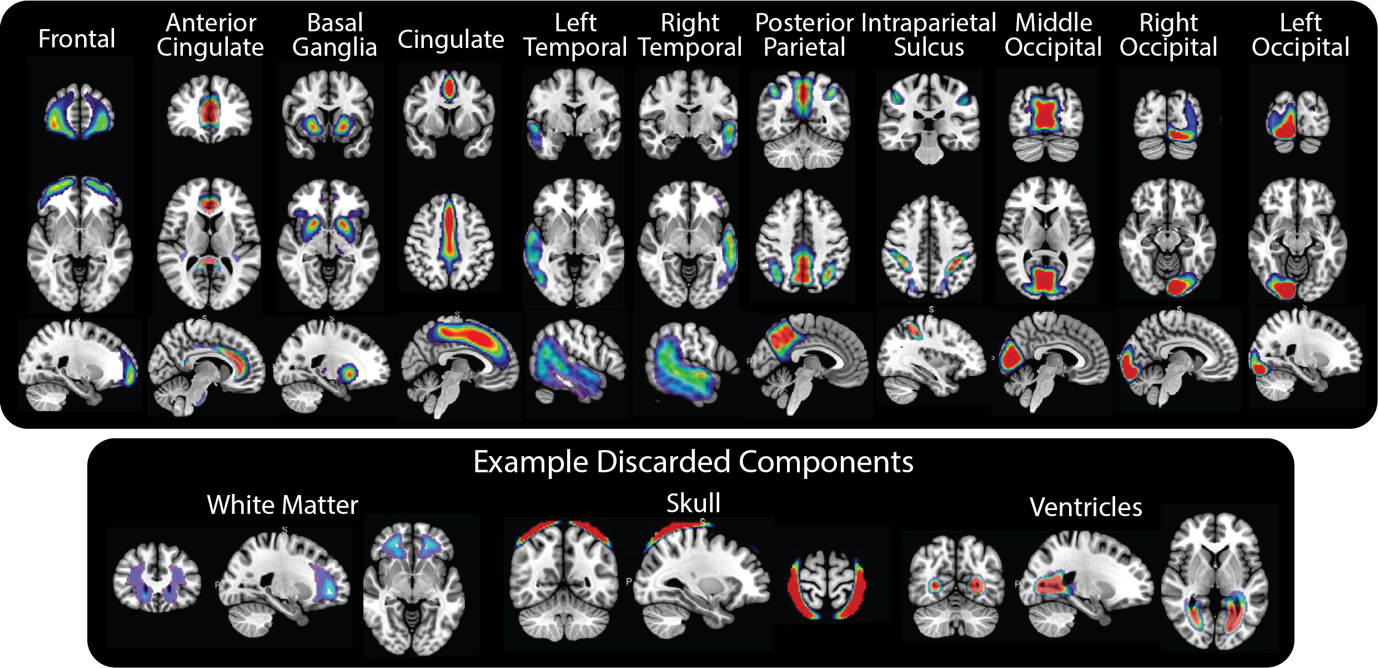

**Supplementary Figure 1.** Group ICA grey matter component loadings projected onto the MNI brain. Loading matrices are estimated on the IDEAS sample and used to derive ICA scores in holdout data sets. Heat map shows the positive tail of the ICA components used to derive the ICA grey matter scores. Top panel shows the 11 components shown were retained from the full 40-dimensional ICA as they capture grey matter Aβ-PET binding. Bottom panel shows three examples of non-grey matter components that were excluded capturing signal from the white matter, skull, and ventricles.

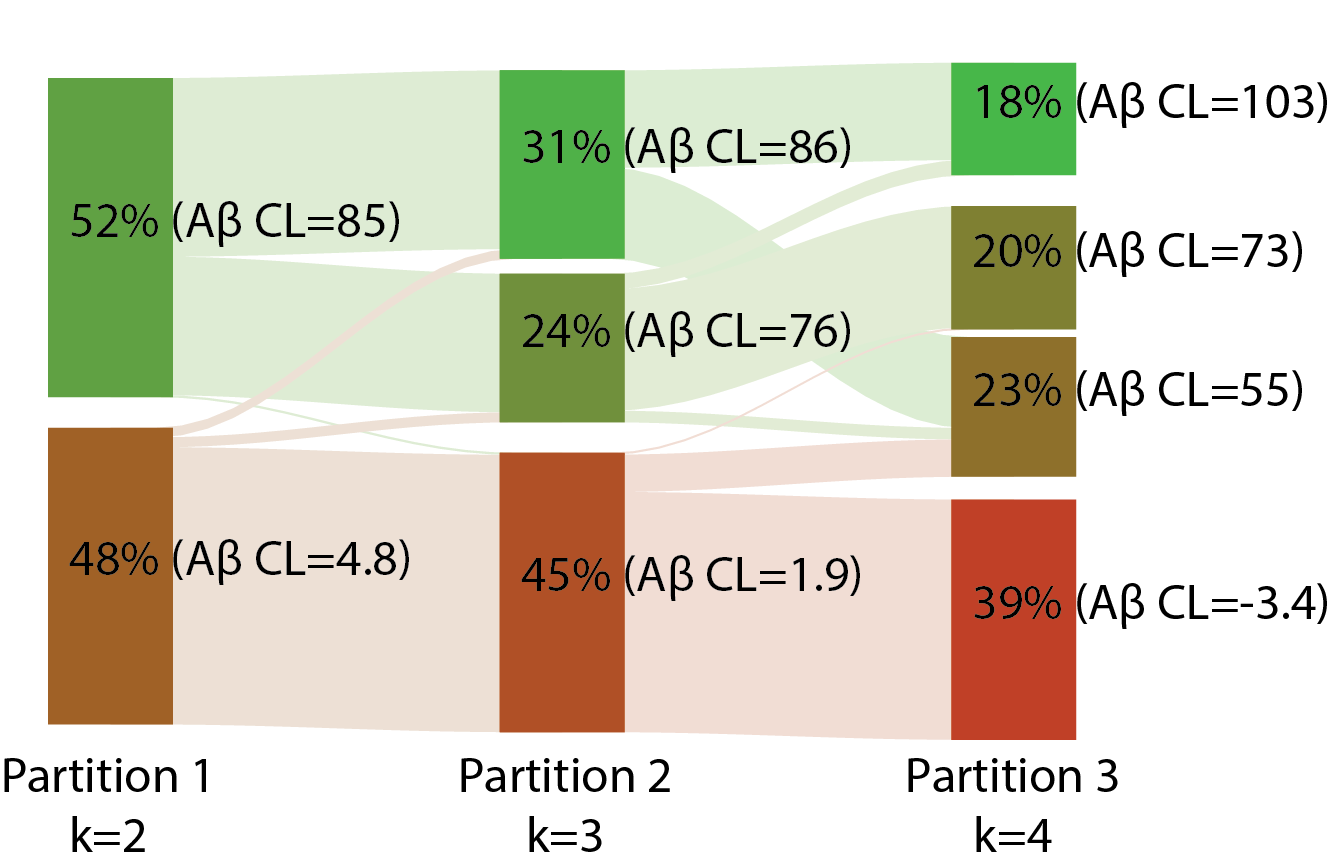

**Supplementary Figure 2.** Alluvial flow diagram showing IDEAS sample partitions at increasing dimensionality of k-means clustering. Partition 3 (k neighbours = 4) shows that one population (mean Aβ CL=73) is almost entirely preserved from Partition 2 (k neighbours = 3). Note that the nested partitioning of clusters is not enforced in k-means, rather the splitting of clusters at higher dimensions is derived by the structure of the data (i.e. Euclidean distances).

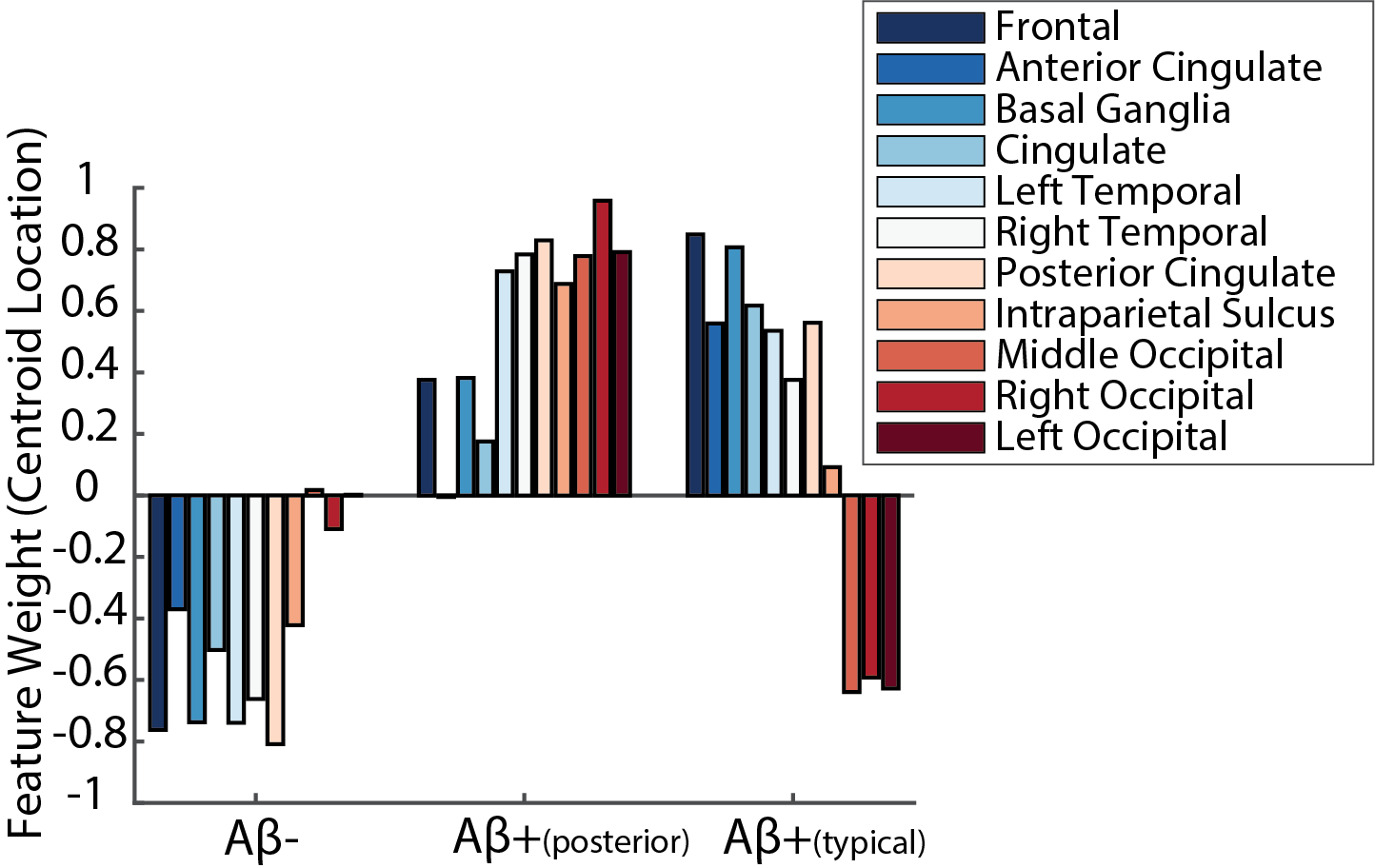

**Supplementary Figure 3.** Centroid locations from the k-means clustering of grey matter Aβ PET components. k-means clustering was run only on the IDEAS sample. Stacked bars represent the centroid of each cluster, values are normalised and thus represent the relative contribution of each grey matter component used as input in clustering.

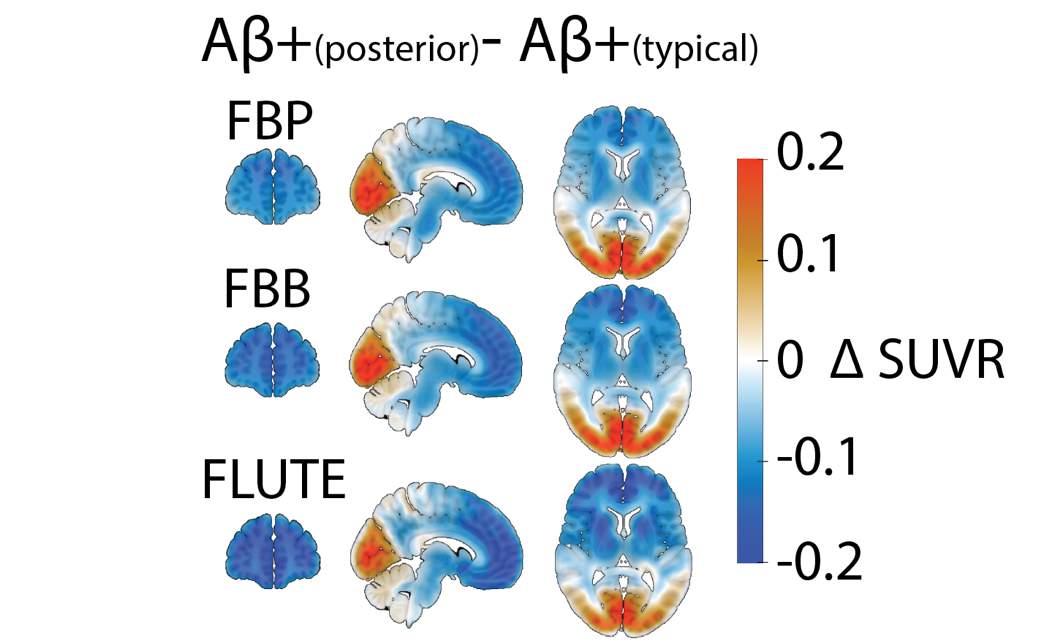

**Supplementary Figure 4.** Difference between average Aβ PET standardised uptake value ratio (SUVR) for Aβ+_(posterior)_ vs. Aβ+_(typical)_ participants from the IDEAS sample separated by Aβ PET tracer: [18]F-Florbetaben (FBB); [18]F-Florbetapir (FBP); [18]F-Flutemetamol (FLUTE).

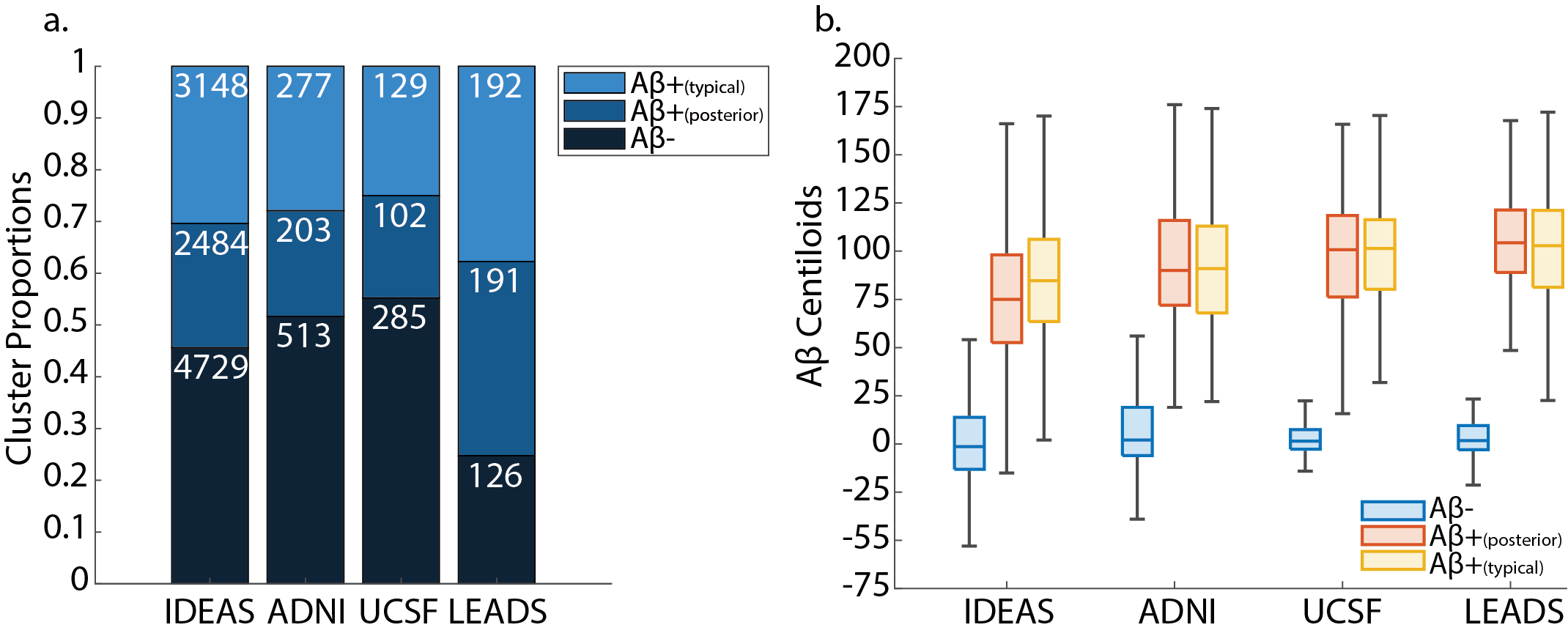

**Supplementary Figure 5. a.** Proportion of participants assigned to each Aβ PET cluster across the four cohorts, white values indicate the total number of patients assigned to each group. b. Distribution of Aβ centiloid (CL) values in each Aβ PET cluster across the four cohorts (Aβ+_(posterior)_ vs. Aβ+_(typical):_ IDEAS mean CL 75.8 vs. 85.6, t(5630)=-11.1, p<0.001; LEADS mean CL 105.4 vs. 101.1, t(377)=1.155, p=0.121; ADNIL mean CL 94.4 vs. 91.6, t(478)=0.948, p=0.344; UCSF mean CL 97.4 vs. 98.9 t(229)=-0.396, p=0.692).

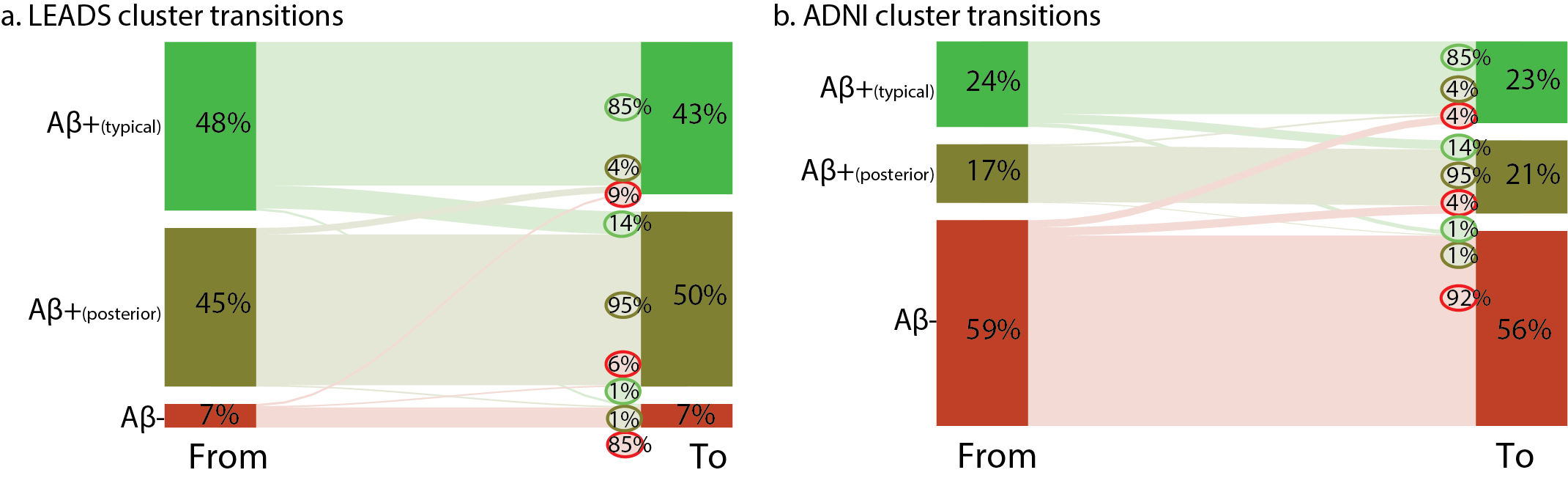

**Supplementary Figure 6.** Alluvial flow diagram showing the temporal evolution of different Aβ clusters in a. LEADS and b. ADNI patients with longitudinal Aβ PET imaging. Percentages at each block indicate the proportion of the entire sample at the first and subsequent sessions. Percentages in the circles at the right block indicate the transition percentages from converting from Aβ- (red); Aβ+_(posterior)_ (dark green); Aβ+_(typical)_ (light green) to Aβ-; Aβ+_(posterior)_; Aβ+_(typical)_. Coloured circles represent the originating cluster and are shown adjacent to the subsequent cluster at follow-up.

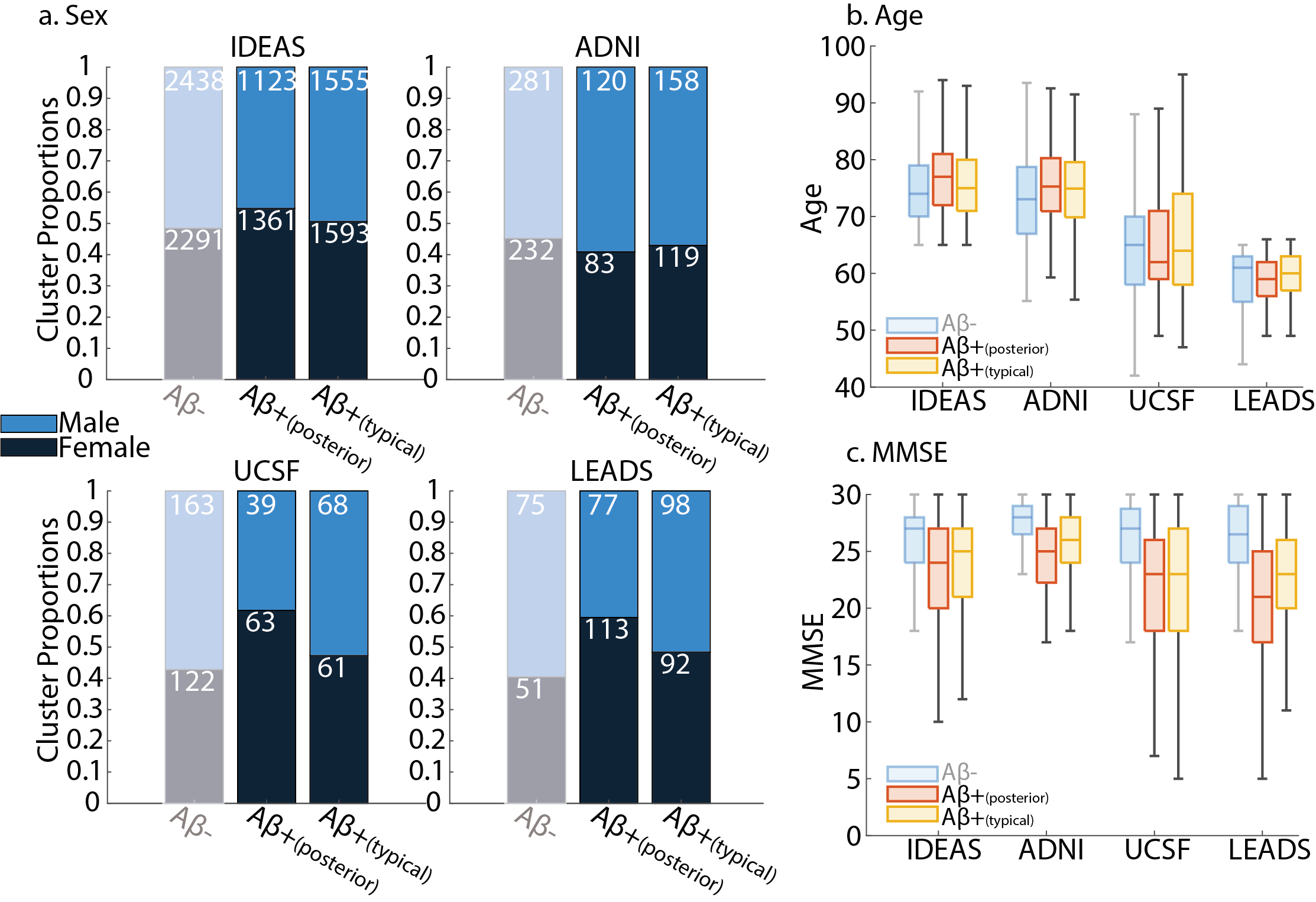

**Supplementary Figure 7.** a. Proportion of males and females assigned to each Aβ PET cluster across the four cohorts (%female Aβ+_(posterior)_ vs. Aβ+_(typical),_ IDEAS: 55% vs. 50%, χ^2^=9.76, p=0.002; LEADS: 59% vs. 48%, χ^2^=4.671, p=0.03; ADNI: 41% vs. 43%, χ^2^=0.2067, p=0.65; UCSF: 62% vs. 47%, χ^2^=4.802, p=0.03).b. Distribution of Age in each Aβ PET cluster across the four cohorts (IDEAS t(5630)=5.9, p<0.001; ADNI t(478)=0.7993, p=0.42; UCSF t(229)=-0.64, p=0.52; LEADS t(378)=-1.592, p=0.11). c. Distribution of mini mental state examination (MMSE) scores in each Aβ PET cluster across the four cohorts (Aβ+_(posterior)_ vs. Aβ+_(typical_: IDEAS t(4337)=-2.263, p<0.001, Cohens D =-0.069; LEADS t(376)=-3.938, p<0.001, Cohens D =-0.404; ADNI t(478)=-3.339, p<0.001, Cohens D =-0.308; UCSF t(222)=-0.51, p=0.61, Cohens D =-0.068). We show as semi-transparent data in each panel the values for the Aβ- cluster.

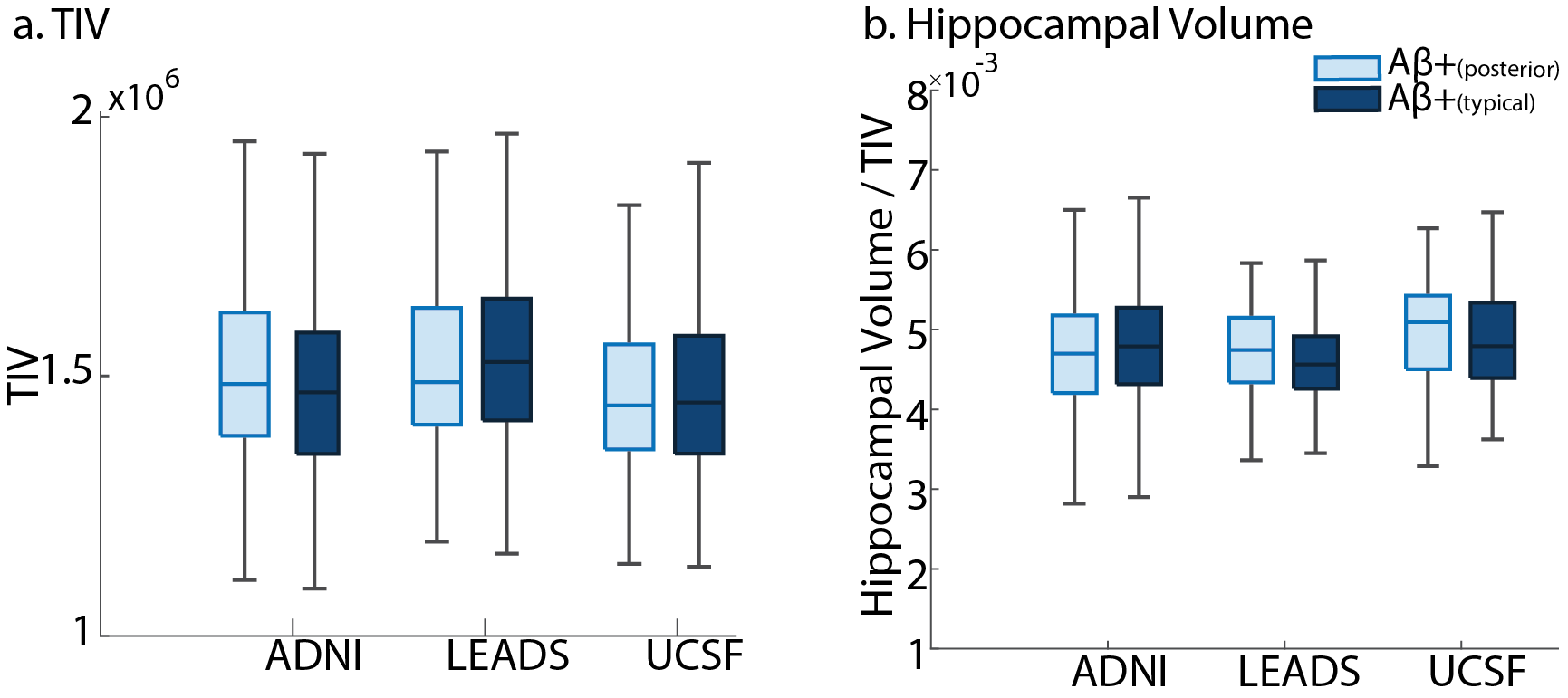

**Supplementary Figure 8** a. Distribution of head size measured by total intracranial volume (TIV) in Aβ+ PET clusters cluster across the cohorts with MRI imaging available (ADNI TIV t(478)=1.2806, p=0.20; UCSF TIV t(227)=0.6059, p=0.55; TIV t(368)=-1.6389, p=0.10). b. Distribution of hippocampal volume normalised by TIV in Aβ+ PET clusters across the cohorts with MRI imaging available (ADNI HV t(478)=0.125, p=0.90; UCSF HV t(227)=0.6908, p=0.49; LEADS HV t(368)=2.664, p=0.008).

|  | **UCSF** | **ADNI** | **Combined** |
| --- | --- | --- | --- |
| **Sample size** | 166 | 49 | 215 |
| **Age mean ± std** | 65±8.9 | 79±7.5 | 68.3±10.5 |
| **Female Sex (%F)** | 78 (47%) | 13 (27%) | 91 (42%) |
| **MCI/Dementia (%Dem)** | 69/97±(58%) | 11/38 (78%) | 80/135 (63%) |
| **MMSE mean ± std** | 22.6±6.4 | 22.6 ±5.6 | 22.6±6.2 |
| **Centiloids mean ± std** | 51.8±53.5 | 71.3±51.8 | 56.2±53.6 |
| **APOE-ε4 (0/1/2) alleles** | 101/50/12 | 21/22/6 | 122/72/18 |
| **Cluster assignment**  **(Aβ-/Aβ+_(posterior)_ /Aβ+_(typical)_)** | 87/38/41 | 13/16/20 | 100/54/61 |

**Supplementary Table 1. Neuropathology sample characteristics.**

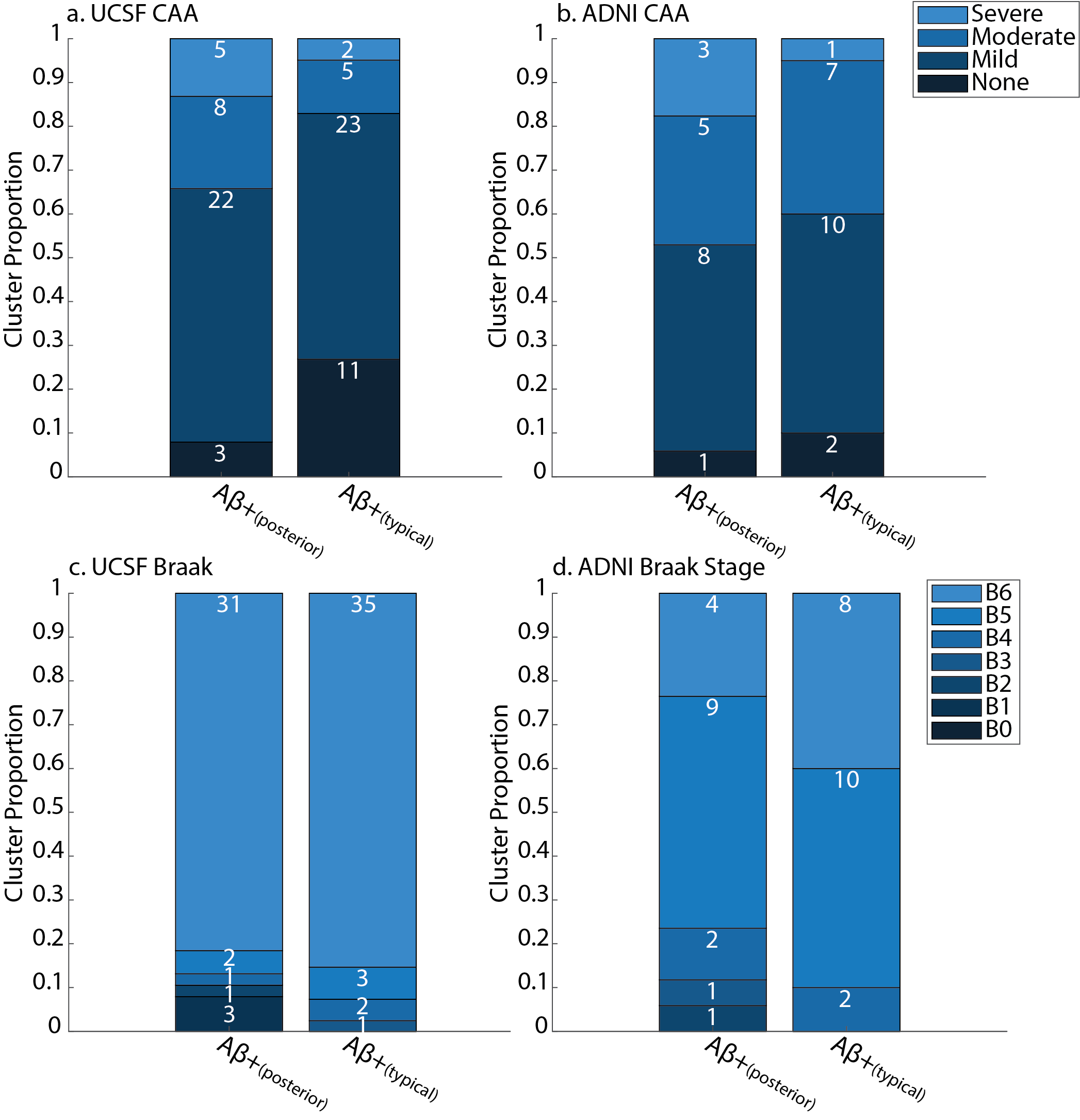

**Supplementary Figure 9** Severity of cerebral amyloid angiopathy (CAA) in each Aβ+ PET cluster in the a. UCSF (UCSF t(77)=2.47, p=0.016 β=0.17) and b. ADNI (ADNI t(35)=0.90, p=0.38 β=0.09) samples separately. Severity of tau pathology assessed using Braak staging in each Aβ+ PET cluster in the c. UCSF (UCSF t(77)=1.40, p=0.164 β=0.07) and d. ADNI (ADNI t(35)=1.66, p=0.11 β=0.15) samples separately. Stacked barcharts represent the proportion of each group, where the white number shows the number of participants represented in each segment.

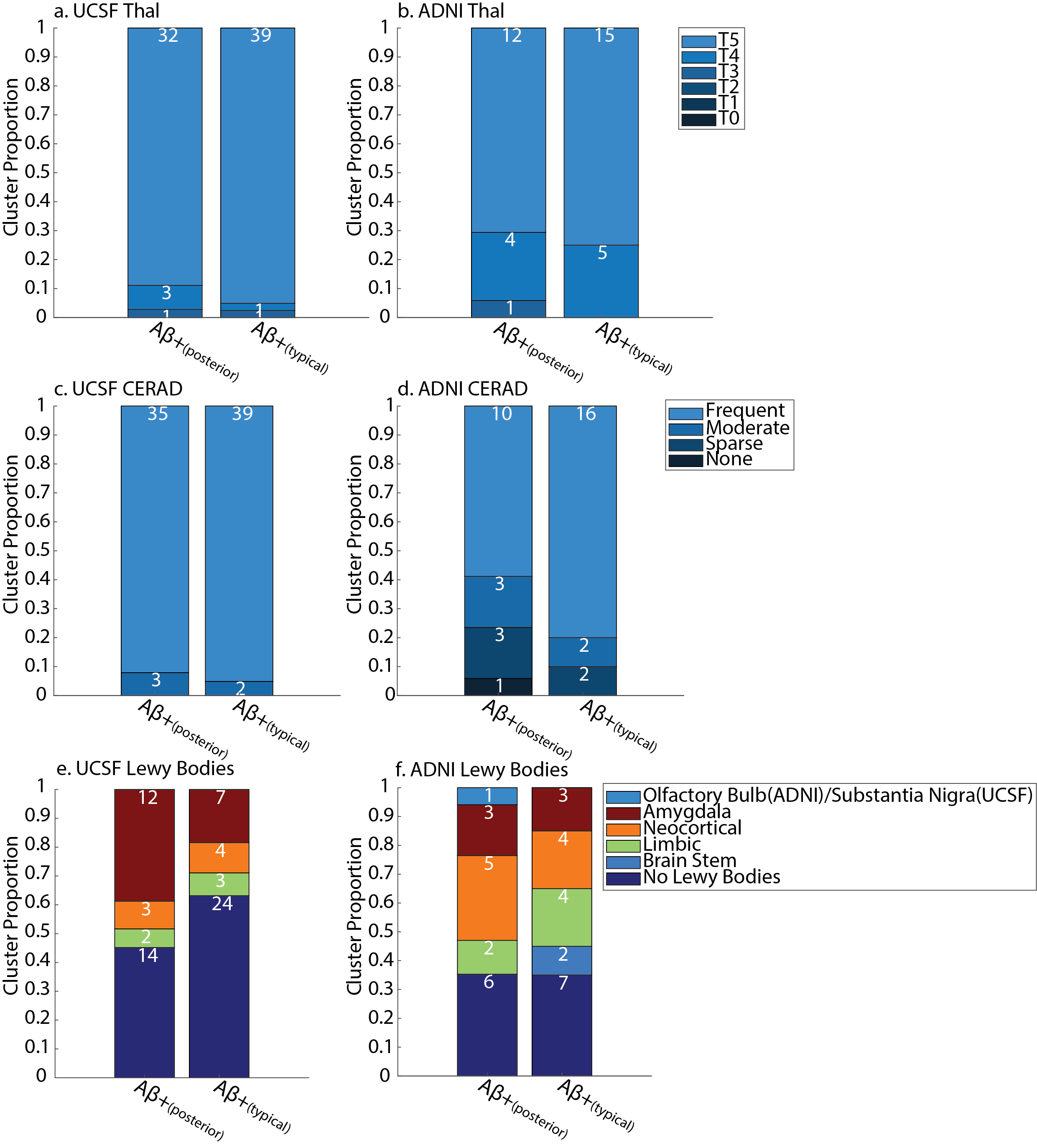
**Supplementary Figure 10** Severity of cerebral amyloid pathology assessed using Thal staging in each Aβ+ PET cluster in the a. UCSF and b. ADNI samples separately. Severity of neuritc plaque density using the Consortium to Establish a Registry for Alzheimer Disease (CERAD) scale in the c. UCSF and d. ADNI samples separately. Distribution of Lewy bodies in the e. UCSF and f. ADNI samples separately. Stacked barcharts represent the proportion of each group, where the white number shows the number of patients represented in each segment.

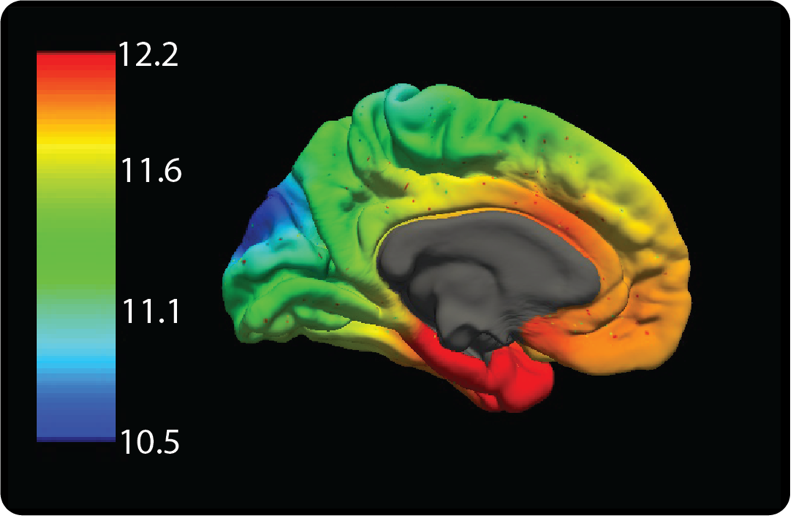

**Supplementary Figure 11** APOE gene expression across the cortex. Colour scale represents log2 mRNA expression intensity. Figure generated online (<https://www.meduniwien.ac.at/neuroimaging/mRNA.html>) as described in Gryglewski et. al. 2018^1^
